# Phosphatase-independent activity of smooth-muscle calcineurin orchestrates a gene expression program leading to hypertension

**DOI:** 10.1101/2023.11.26.568733

**Authors:** Paula Sofía Yunes-Leites, Yilin Sun, Sara Martínez-Martínez, Álvaro Alfayate, Marta Toral, Ángel Colmenar, Ana Isabel Torralbo, Dolores López-Maderuelo, Sergio Mateos-García, David N. Cornfield, Jesús Vázquez, Juan Miguel Redondo, Miguel R. Campanero

## Abstract

**Objective:** Angiotensin-II (Ang-II) drives pathological vascular wall remodeling in hypertension and abdominal aortic aneurysm (AAA). Previous studies showed that the phosphatase activity of calcineurin (Cn) mediates Ang-II-induced AAA, but the cell type involved in the action of Cn in AAA formation remained unknown.

**Methods:** Smooth muscle cell (SMC)-specific and endothelial cell (EC)-specific Cn-deficient mice (*SM-Cn^-/-^* and *EC-Cn^-/-^* mice, respectively) were created and assessed for Ang-II-induced AAA formation and hypertension vs controls. Osmotic minipumps were used to administer Ang-II and cyclosporine A (CsA), a pharmaceutical Cn inhibitor. AAA formation and hypertension were monitored by ultrasonography, arterial blood pressure monitoring, and histological analysis. Deep RNA sequencing was used to identify the Ang-II-regulated transcriptome sensitive to Cn deletion or pharmacological inhibition. Arterial and SMC contractility were also assessed.

**Results:** Cn expressed in SMCs, but not ECs, was required for Ang-II-induced AAA. Unexpectedly, SMC Cn played a structural role in the early onset and maintenance of Ang-II-induced hypertension independently of Cn phosphatase activity. Nearly 90% of the genes regulated by Ang-II in the aorta required Cn expression in SMCs. Cn orchestrated, independently of its enzymatic activity, the induction by Ang-II of a gene expression program closely related to SMC contractility and hypertension. Cn deletion in SMCs, but not its pharmacological inhibition, impaired the regulation of arterial contractility. Among the genes whose regulation by Ang-II required Cn expression but not its phosphatase activity, we discovered that *Serpine1* was critical for Ang-II-induced hypertension. Indeed, pharmacological inhibition of PAI-1, the protein encoded by *Serpine1*, impaired SMCs contractility and readily regressed hypertension.

**Conclusions:** Whereas the phosphatase activity of Cn mediates Ang-II-induced AAA, a phosphatase-independent action of SMC Cn mediates hypertension by orchestrating a gene expression program closely related to contractility and blood pressure regulation. Our results urge the evaluation of PAI-1 as a candidate therapeutic target for hypertension.

## INTRODUCTION

Arteries are complex structures composed of 3 distinct layers—the intima, media, and adventitia—each of which plays a unique physiological role. The intima is a single layer of endothelial cells (ECs), whereas the adventitia is a composite of various cell types and extracellular matrix (ECM) components. The media comprises multiple layers of ECM and vascular smooth muscle cells (VSMCs), whose contractile ability provides the basis of the overall contractility of the vessel. However, changes in hemodynamic forces or molecular signaling pathways can lead to pathological vascular wall remodeling (VWR), resulting in destabilization of the arterial wall and potentially leading to outcomes such as aortic aneurysm (AA) or arterial hypertension.

AA is a progressive and abnormal enlargement and weakening of the aortic wall that can eventually result in aortic dissection (AD) or rupture, accounting for 1–2% of all deaths in industrialized countries. Before dissection or rupture, AA is often clinically silent, and there is therefore a clear need for improved techniques for early diagnosis. Key features of AA include upregulation of VSMC proliferation, migration, and apoptosis and the repression in these cells of genes related to quiescence and contractility ^1^.

Hypertension is one of the most common chronic diseases, affecting more than 1 billion people worldwide. Hypertension is considered a multiorgan outcome, with the underlying cause identified in only 5% of patients ^2^. Blood pressure is determined by the contractile state of the arteries, which can be dysregulated by a variety of VSMC-mediated processes, including hyperplasia, hyperproliferation, and increased contractility ^3,4^.

VWR is triggered by angiotensin-II (Ang-II), a key effector of the renin-angiotensin system. In abdominal AA (AAA), Ang-II activates pro-mitogenic and pro-apoptotic pathways, increases ECM deposition, and induces cell migration and inflammation ^5,6^. While the importance of Ang-II in VWR is broadly accepted, the underlying molecular mechanisms are incompletely understood. Ang-II exerts its pathological effects by binding to its type 1 receptor (AT1R), which activates a variety of signaling pathways, including those mediated by increased cytosolic [Ca^2+^], activation of the PKC-dependent tyrosine kinase cascade, NADPH-dependent ROS production, and activation of JAK/STAT, AKT, and MAPK kinases ^6^.

One of the key effectors activated by the increase in cytosolic [Ca^2+^] is the ubiquitously expressed Ca^2+^/calmodulin-dependent serine/threonine phosphatase calcineurin (Cn). Cn, comprising a catalytic subunit (CnA) and a regulatory subunit (CnB), plays a key role in the development of pathological VWR by mediating Ang-II-induced VSMC migration ^7^ and senescence ^8^, regulating endothelial nitric oxide synthase activity and endothelial barrier function in ECs ^9,10^, and mediating fibroblast migration and collagen secretion ^11^. Inhibition of Cn activity with cyclosporine A (CsA) diminishes Ang-II-triggered neointima formation and AAA development, which is also inhibited by lentiviral expression of Cn-blocking peptides ^7,12^. Despite the importance of Cn in pathological VWR, knowledge is lacking about the relative contribution of Cn expressed in the different vascular cell types to the development of vascular disease.

## METHODS

### Mouse strains

To delete Cn specifically in smooth muscle cells or endothelial cells, we crossed Cnb1^flox/flox^ (Cn^f/f^) mice ^13^ with either *Cdh5-Cre^ERT2^* transgenic mice ^14^ to obtain EC-Cn^f/f^ mice or with *Myh11-Cre^ERT2^* transgenic mice ^15^ to obtain SM-Cn^f/f^ mice. All mouse strains were backcrossed with C57BL/6J mice for more than 8 generations to obtain a homogeneous genetic background and were maintained in homozygosity for the *CnB1*-floxed locus and in hemizygosity for the Cre^ERT2^ transgene in the case of EC-Cn^f/f^ and SM-Cn^f/f^ mice. All mice were genotyped by PCR of genomic DNA obtained from tail samples (genotyping PCR). In some experiments, genomic DNA samples from tamoxifen-treated mice were analyzed by agarose gel electrophoresis to detect the Cre-loxP recombination (recombination PCR). Primer sequences used for genotyping were: *CnB1-*floxed mice (5’-CCCTAGGCACTGTTCATGGT-3’, 5’-GCCACAGATTAGCCTCGTGT-3’, 5’-CACATTCACCCACAATTGC-3’), *Myh11-Cre^ERT2^* mice (5’-TGACCCCATCTCTTCACTCC-3’, 5’-AACTCCACGACCACCTCATC-3’, 5’-AGTCCCTCACATCCTCAGGTT-3’), and *Cdh5-Cre^ERT2^* mice (5’-GCCTGCATTACCGGTCGATGCAACGA-3’, 5’-GTGGCAGATGGCGCGGCAACACCATT-3’).

### Animal procedures

Animal procedures were approved by the CNIC Ethics Committee and by the Madrid regional authorities (ref. PROEX 80/16 and PROEX 182.1/20) and conformed to European regulations regarding animal care (Directive EU 2010/63EU and Recommendation 2007/526/EC). All mice were housed in the CNIC specific pathogen-free animal facility with controlled temperature, humidity, and light-dark cycles. Sacrifice of mice was done by CO2 inhalation.

*Cnb1*^f/f^-Cre^ERT2-neg^ together with *Cnb1*^wt/wt^;*Cdh5*-Cre^ERT2^ or *Cnb1*^wt/wt^;*Myh11*-Cre^ERT2^, were used as controls (Cn-Ctl) for EC-KO or SM-KO, respectively. For inducible and conditional gene deletion, 10–12-week-old Cn-Ctl, SM-Cn^f/f^, and EC-Cn^f/f^ male mice were given daily 1 mg i.p. injections of tamoxifen (Sigma-Aldrich) on 5 consecutive days, generating SM-Cn^-/-^ and EC-Cn^-/-^ mice. Cn-Ctl, SM-Cn^-/-,^ and EC-Cn^-/-^ male mice were used after 6 weeks of tamoxifen administration.

Ang-II (Sigma-Aldrich), CsA (Novartis), NE (Sigma-Aldrich), and TM5441 (MedChemExpress) were administered at 1 μg/kg/min, 5mg/kg/d, 10 mg/kg/d, and 20mg/kg/d, respectively, using subcutaneous osmotic minipumps (Alzet Corp, model 2001 or 2004). *N_ω_*-nitro-L-arginine methylester hydrochloride (L-NAME; 0.5 g/l; Sigma-Aldrich, N5751) was administered in drinking water every 3 days. For minipump implantation, all mice were isoflurane-anesthetized (3-5% isoflurane for induction, 1-2% isoflurane for maintenance), and a small surgical pocket was made in the dorsal skin, where the minipump was implanted. Mice undergoing the same surgical procedure but without minipump implantation were used as controls for the Cn-Ctl, SM-Cn^-/-^, EC-Cn^-/-^, and CsA-treated groups.

For the AAA model, Cn-Ctl, SM-Cn^-/-^, and EC-Cn^-/-^ male mice were fed a HFD (EF M HF coconut oil, +7.5 g/kg Cholesterol Experimental diet, 38% chow, 10 mm; S9167-E011, SSNIFF) for 12 consecutive weeks. In the first week of feeding, tamoxifen was injected to induce deletion in the Cre^ERT2^-expressing mice. After 12 weeks of the HFD, this treatment was combined with the systemic infusion (1 µg/kg/min) of Ang-II (Sigma-Aldrich) with osmotic minipumps (Alzet Corp) ^16^.

For the SM-Cn^-/-^ transcriptome analysis, short-term Ang-II infusion and mouse selection were performed as follows: male mice were treated with tamoxifen and, after 6 weeks, baseline BP measurements were performed. Osmotic minipumps with Ang-II (1 µg/kg/min) were implanted in Cn-Ctl and SM-Cn^-/-^ mice in parallel, and BP was measured after 2h and 24h. Treated Cn-Ctl mice with systolic BP >130 mmHg were considered valid for analysis and were euthanized; all the SM-Cn^-/-^ mice treated in parallel were selected for analysis, since none of them showed increased BP after Ang-II infusion. For the CsA transcriptome analysis, osmotic minipumps with CsA (CsA group) were implanted in WT C57BL/6J male mice. Minipumps were not implanted in control mice at this stage. After 5 days, baseline BP measurements were taken. Ang-II osmotic minipumps (1 µg/kg/min) were implanted in CsA and control mice in parallel, and BP was measured after 2h and 24h. Ang-II-treated CsA and control mice with a systolic BP >130 mmHg were considered valid for analysis and euthanized. If we could not confirm *CnB1* deletion in any SM-Cn^-/-^ mouse or inhibition of NFAT dephosphorylation in the thymus of any CsA-treated mouse, those mice were removed from the experiment. Experiments for validation by RT-qPCR and protein expression by western blotting were performed in a similar way.

### Blood pressure measurements

Arterial blood pressure (BP) was measured by the mouse tail-cuff method using the automated BP-2000 Blood Pressure Analysis System (Visitech Systems, Apex, NC, USA). In brief, mice were trained for BP measurements every day for one week. After training, BP was measured one day before treatment to determine the baseline BP values in each mouse cohort. Measurements were repeated several times during experiments. BP measurements were recorded in mice located in a tail-cuff restrainer over a warmed surface (37°C). Fifteen consecutive systolic and diastolic BP measurements were made, and the last 10 readings per mouse were recorded and averaged.

### In vivo ultrasound imaging

Ultrasound images were obtained from isoflurane-anesthetized mice (1.5-2% isoflurane) by high-frequency ultrasound with a VEVO 2100 echography device (VisualSonics, Toronto, Canada) at 30-micron resolution. Echocardiography was used to measure diastolic maximal internal aortic diameters and interventricular septum thickness in the parasternal long-axis view in the two-dimensional guided M-mode. Measurements were taken before treatment initiation to determine the baseline diameters and were repeated several times during the experiment. At endpoint, aortas were classified as AAA when they had increased in diameter by 50% or more ^17,18^, as dilated when the increase was 20-50%, and as normal when the increase was 20% or less, relative to the initial measurement in all cases.

### Lentivirus production and infection

Lentivirus expressing Cre recombinase, GFP protein, LxVP, and LxVPmut peptides have been described previously ^7,19^. Pseudo-typed lentivirus production and titration was as described previously ^19,20^. Briefly, HEK-293T cells were transiently transfected using the calcium phosphate method, and culture supernatant was concentrated by ultracentrifugation (2h at 26,000rpm; Ultraclear Tubes; SW28 rotor and Optima L-100 XP Ultracentrifuge; Beckman). Viruses were suspended in cold sterile PBS and titrated by transduction of Jurkat cells for 48h. Transduction efficiency (GFP-expressing cells) and cell death (propidium iodide staining) were quantified by flow cytometry (BD FACSCanto Flow Cytometer and FlowJo^TM^ v10.8 software). The HEK-293T and Jurkat cell lines were purchased from ATCC. All cells were mycoplasma-negative.

### Lentiviral transduction of aortas in vivo

WT male mice were anesthetized by intraperitoneal injection of ketamine (120 mg/kg) and xylazine (16 mg/kg), and a small incision was made to expose the jugular vein ^7,20^. Virus solution (200 μl, 10^9^ particles/ml in PBS) was inoculated directly into the jugular vein 6 weeks before Ang-II minipump implantation. Transduction efficiency was analyzed postmortem in aortic samples by GFP immunohistochemistry.

### Cell procedures

Primary mouse vascular smooth muscle cells (VSMCs) were isolated and grown as described ^21^. In brief, aortas from Cn^f/f^ male mice were dissected, and the adventitia was removed with forceps. Aortic tissue was cut into small rings and digested with 1 mg/ml collagenase type II (Worthington) and 0.5 mg/ml elastase (Worthington) in DMEM (Gibco) until single cells were observed under the microscope. The enzymatic reaction was stopped by the addition of DMEM supplemented with bovine serum. For primary mouse aortic endothelial cells (mAECs), digested tissue was incubated with anti-ICAM-2 (BD Biosciences, 553326) conjugated to magnetic beads (Dynal Thermofisher, 11035). Isolated cells were cultured, and the remaining tissue was removed in the following days.

VSMCs were cultured in DMEM (Gibco) supplemented with 20% FBS (Gibco) and antibiotics. mAECs were cultured in DMEM F-12 (BioWhittaker BE, 12-719F) supplemented with 20% FBS, 100 mg/ml heparin (Sigma-Aldrich), 5 mg/ml endothelial cell growth factor (Gibco), and antibiotics on culture plates pre-covered with 0.5% gelatin and 100mg/ml collagen type I (Sigma-Aldrich, C2124-50ML). EC purity was >80% after 1 week of culture. All experiments were performed in passages 3-5, and all cells were mycoplasma-negative.

VSMCs were infected with specific lentiviruses (LVs) for 5h at a multiplicity of infection (MOI) = 10. The medium was then replaced with fresh supplemented DMEM. Seven days after the transduction, infection efficiency was evaluated and experiments were performed. In the case of Cre-recombinase-expressing LVs, infection was confirmed by immunoblot detection of Cn deletion. In the case of LxVP-expressing LVs, infection was confirmed by RT-qPCR detection of GFP.

For collagen contraction assays, 150 µl aliquots of PureColTM EZ Gel Solution (Sigma-Aldrich, 5074) were polymerized for 1h at 37°C in p48 wells, and 8×10^4^ VSMCs were seeded on each gel. Cells were left to adhere to the gels for 5h and then starved for 72h. The gel–cell composites were then gently detached from the plates and incubated in DMEM supplemented with 3% FBS for 24h-48h. When indicated, cells were pretreated with 50 µM TM5441 (MedChemExpress) for 2h. At the end of the experiment, gels were washed in PBS and fixed with 3% paraformaldehyde (PFA) for 15 minutes. Gel images were taken with a Nikon SMZ800 scoop camera (0.8x objective). Each condition was replicated in 3-4 wells, and each condition was replicated with different batches of VSMCs. Gel contraction was quantified with ImageJ software as follows: each gel area was quantified and normalized to the average of all the areas in a single experiment; the means of the normalized areas for each condition are presented.

### Myography

Rings (3 mm long) from control (Cn-Ctl or WT) and SM-Cn^-/-^ male mouse thoracic descending aortas and superior mesenteric resistance arteries (MRA) were excised and placed on a wire myograph (model 610M, Danish Myo Technology, Aarhus, Denmark) for the measurement of isometric tension. The organ chamber was filled with Krebs solution (composition in mM: NaCl 118, KCl 4.75, NaHCO_3_ 25, MgSO_4_ 1.2, CaCl_2_ 2, KH_2_PO_4_ 1.2, and glucose 11) and infused with carbogen (5% CO_2_ and 95% O_2_, pH ∼7.4), at 37°C. When indicated, CsA (200 ng/ml) was added to WT arteries at the time of removal from the animal and was maintained during ring excision and throughout the experiment in the chamber. Aortic and mesenteric resistant artery (MRA) rings were stretched to tensions of 5 mN and 3 mN, respectively, and equilibrated for 30 min. Tension characteristics were obtained with the myograph software (LabChart). Aortic and MRA rings were initially stimulated with 80 mM and 50 mM KCl, respectively, followed by determination of KCl concentration-response curves (1 mM to 100 mM). Vasoconstriction measurements are presented as the force (mN) exerted by the rings.

### Aortic histology

After sacrifice of mice by CO_2_ inhalation, aortas were perfused with saline, isolated, and fixed in 4% paraformaldehyde overnight at 4°C. Paraffin cross sections (5 μm) from fixed aortas were stained with Masson’s trichrome (Masson), Hematoxylin and Eosin (HE), or Elastic Verhoeff–Van Gieson (EVG) stain. Medial wall thickness was determined from HE staining by quantifying the area of the media in 4 non-consecutive sections containing complete aortic cross sections using the NDP.view2 Image viewing software.

For immunohistochemistry, deparaffinized sections (5 µm) were rehydrated, boiled to retrieve antigens (10 mM citrate buffer, pH 6), and blocked for 1h with 5% goat serum plus 2% BSA in PBS. Samples were incubated with anti-GFP primary antibody (Invitrogen, A11122, 1:200) overnight at 4° and then with a biotinylated secondary antibody for 1h at room temperature. Signal was detected with the DAB (3,3′-Diaminobenzidine) reaction (Vector Laboratories) for a maximum 7-minute reaction time. Sections were then counterstained with hematoxylin, dehydrated, and mounted with DPX (CasaÁlvarez).

*In situ* southwestern histochemistry was performed as previously described ^7^. Briefly, 5 μm aortic cross sections were deparaffinized and rehydrated. Sections were fixed with 0.5% paraformaldehyde for 25 min at 28°C, and endogenous alkaline phosphatase was quenched with 5 mM levamisole (L9756, Sigma-Aldrich). Preparations were then digested with 0.5% pepsin A (P7000, Sigma-Aldrich) in 1 N HCl for 35 min at 37°C and washed twice with HEPES/BSA buffer (10 mM HEPES, 40 mM NaCl, 10 mM MgCl_2_, 1 mM dithiothreitol [DTT], 1 mM EDTA, and 0.25% BSA, pH 7.4). Sections were incubated with 0.1 mg/ml DNase I in HEPES/BSA buffer for 30 min at 30°C. Activated NFAT proteins were probed with the synthetic sense DNA sequence 5’-GATCGCCCAAAGAGGAAAATTTGTTTCATACAG-3’. This sequence contains a composite site comprising the NFAT and AP1 sites from the mouse *IL-2* promoter ^22^. The NFAT probe was labeled with digoxigenin (DIG) (Sigma-Aldrich) and diluted to 25 pM in HEPES/BSA buffer containing 0.5 mg/ml poly (dI-dC) (P4929, Sigma-Aldrich). Absence of probe was used as negative control. After overnight incubation at 37°C in a humidified chamber, sections were washed twice with buffer 1 (0.1 M maleic acid, 0.15 M NaCl, 0.03% Tween-20, pH 7.5) and incubated for 1 h in blocking solution (0.1% SSC and 0.1% SDS diluted 1:10 in buffer 1 without Tween). The preparations were washed with buffer 1 and incubated for 2h at 37°C with an anti-digoxigenin antibody conjugated to alkaline phosphatase (1:200 in blocking solution; 11093274910, Roche). Next, sections were washed in buffer 1 and in buffer 3 (Tris HCl 0.1 M, 0.1 M NaCl, and 50 mM MgCl_2_, pH 9.5) at room temperature. Bound alkaline phosphatase was visualized with nitroblue tetrazolium chloride and 5-bromo-4-chloro-3-indolyl-phosphate (NBT/BCIP; 11681451001, Roche). The reaction was stopped by incubation in buffer 4 (10 mM Tris and 1 mM EDTA, pH 8.0), and sections were dehydrated through a graded ethanol series and mounted in 50% glycerol in PBS.

All bright-field images were obtained with a Leica DM2500 microscope with 20x and 40x HCX PL Fluotar objectives and with Leica Application Suite V3.5.0 software. Southwestern histochemistry images were analyzed and quantified with ImageJ software as follows: after color deconvolution by fast red blue DAB, the blue channel was selected (color 2). The same threshold was selected in all images, and only the medial aortic region was selected as a region of interest (ROI) and measured. Raw data are presented. Layouts were obtained in Adobe Illustrator.

### Immunoblotting

Mouse samples were obtained by dissection and, in the case of aortas, perivascular tissue and adventitia were removed with forceps. The tissue was frozen in liquid nitrogen and then homogenized using a mortar. Cultured cells were washed with ice-cold PBS and frozen at-80°C. Protein extracts from tissue samples and cells were obtained in ice-cold high-salt lysis buffer (20 mM Tris HCl pH 7.5, 5 mM MgCl_2_, 50 mM NaF, 10 mM EDTA, 0.5 M NaCl, 1% Triton X-100) for the detection of CnB, CnA, tubulin, α-SMA, Cd31, and NFATc3, and in RIPA buffer (150mM NaCl, 10mM Tris HCl pH8, 1% Triton X-100, 0.1% SDS, 1% sodium deoxycholate and 5mM EDTA) for the detection of PAI-1 and tubulin. Both lysis buffers were supplemented with protease, phosphatase, and kinase inhibitors. Homogenates were centrifuged for 15 min at 16000 x g at 4 °C, and proteins were collected in the supernatants. Protein was quantified with the Lowry assay kit. Protein samples (30 μg) were separated under reducing conditions on SDS-polyacrylamide gels and transferred to nitrocellulose membranes. Proteins were detected with the following primary antibodies: mouse monoclonal anti-CnB (Sigma, 0581, 1:1000), rabbit polyclonal anti-CnA (Merck Millipore, 07-1491, 1:2000), mouse monoclonal anti-tubulin (Sigma, T6074, 1:40000), rabbit polyclonal anti-Cd31 (Abcam, ab28364, 1:1000), mouse monoclonal anti α-SMA (Sigma, A5228; 1:5000), rabbit polyclonal anti-NFATc3 (M-75) (SCB, sc-8321, 1:1000), and rabbit monoclonal anti-PAI-1 (Abcam, ab222754, 1:500). After washing and incubation with the appropriate specific HRP-conjugated secondary antibodies, immunoreactive bands were visualized using the enhanced chemiluminescence (ECL) detection reagent (GE Healthcare, Chicago, IL, USA, RPN2106) or acquired in the ImageQuant LAS 4000 device using the Immobilon Forte Western HRP substrate (WBLUF0100, Millipore). Quantification of protein band intensity was performed with ImageJ using Tubulin band intensity for normalization.

### RT and quantitative PCR

Aortas were extracted after perfusion with 5 ml saline solution, and the adventitia layer was discarded. Frozen tissue was homogenized using a mortar and an automatic bead homogenizer (MagNA Lyzer). Total RNA was isolated with TRIZOL (Life Technologies). Total RNA (1 μg) was reverse transcribed at 37°C for 50 min in a 20 µl reaction mix containing 200U Moloney murine leukemia virus (MMLV) reverse transcriptase (Life Technologies), 100 ng random primers, and 40U RNase Inhibitor (Life Technologies). The resulting cDNA was analyzed by quantitative real-time PCR (qPCR) in an AB7900 sequence detection PCR system (Applied Biosystem, Foster City, CA, USA) using SYBR Green PCR or TaqMan Universal PCR master mixes (Go Taq Probe PCR Master Mix; Promega, A600A or A610A, respectively). The following TaqMan probes were used: *Ptgs2* (Mm00478374_m1) and *Hprt1* (Mm00446968_m1). The specific probes used for SYBR Green detection were as follows: *Gja1* (5’-CTTGGGGTGATGAACAGT-3’; 5’-TGAGCCAAGTACAGGAGT-3’ ^23^), *Hprt* (5’-GCTGGTGAAAAGGACCTCT-3’; 5’-CACAGGACTAGAACACCTGC-3’), *Ier3* (5’-GCGCGTTTGAACACTTCTCG-3’; 5’-TGGCGCCGGACCACTC-3’), *Rcan1-4* (5’-TTGTGTGGCAAACGATGATGT-3’; 5’-CCCAGGAACTCGGTCTTGT-3’), *Serpine1* (5’-GAGGTAAACGAGAGCGGCA-3’; 5’-AAGAGGATTGTCTCTGTCGGG-3’). Probe specificity was checked by post-amplification melting-curve analysis in the case of SYBR amplification; for each reaction, only one Tm peak was produced. The amount of target mRNA in samples was estimated by the 2^-ΔCT^ relative quantification method, using *Hprt* for normalization and represented in graphs. Each sample represented belongs to one mouse aorta.

### RNA sequencing

Aortas were isolated after perfusion with saline, and the perivascular tissue and the adventitia layer were removed with forceps. A piece of aortic tissue was used for PCR confirmation of *CnB1* recombination, and the thymus was isolated for protein extraction to check the NFAT phosphorylation state, when appropriate. Aortic tissue was frozen in liquid nitrogen and homogenized with lysis buffer (Qiagen, 64204) using an automatic bead homogenizer (MagNA Lyser; Roche). Total RNA was isolated with the RNeasy MinElute Cleanup Kit (QIAGEN, 74204). Each triplicate was a pool of 3 aortas (total, 9 aortas per condition). RNA quality and purity was analyzed by Nanodrop and Bioanalyzer (Agilent 2100 Bioanalyzer RNA NanoChip). Only RNA samples with appropriate absorbance ratios were used (260/280 nm > 2.0; 230/260 nm 2.0–2.2); RNA samples with a RIN (RNA Integrity number-Bioanalyzer) below 7 were discarded. For each condition, 1.5 μg of RNA was retro-transcribed with oligo dT priming, and the cDNA obtained was used to generate the transcriptome library. The library was sequenced using Illumina technology (Illumina Genome Analyzer IIx System). Library generation and sequencing were performed by the CNIC Genomics Unit under the supervision of Dr. Ana Dopazo. Reads were aligned first using BoWTie version 0.12.7 and then mapped to the latest Ensembl transcript set using RSEM v1.16. One replicate of the untreated Cn^f/f^ group showed a high deviation from the other 2 replicates in PCA analysis and was therefore discarded. Genes showing altered expression with an adjusted p < 0.05 were considered differentially expressed genes (DEGs). Only protein-coding genes were included in further analysis.

Venn diagrams were generated with the online software Venny 2.1.0 (https://bioinfogp.cnb.csic.es/tools/venny/index.html), and the DEGs represented in each diagram are indicated in the figure and figure legend. Heatmaps were generated using R library ’stats’ ^24^ and ’gplots’ ^25^ to represent mean normalized mRNA expression of the DEGs regulated by Ang-II in Cn^f/f^, SM-Cn^-/-^, WT, and CsA-pretreated mice vs the untreated condition (FC>±1.5). Mean normalized mRNA expression was obtained from triplicates of each condition. The hypertension score was extracted automatically by an exact pattern match using the gen symbol as the identifier. The processed database was the Harmonizome dataset collection ^26^ for the curated CTD Gene-Disease Associations related to the term ’Hypertension’, where the standardized value was renamed the hypertension score (HS).

### Functional enrichment analysis

Gene Ontology (GO) enrichment analysis of DEGs was performed using g:Profiler software ^27^. The analysis mainly focused on gen function data sources related to cellular components (GO:CC). We were able to map genes to statistically significant enriched terms. The statistical domain scope was only applied to annotated genes, adjusting the significance threshold by multiple testing (using the g:Profiler tailor-made algorithm g:SCS), set to a value p<0.05. This approach has produced more accurate results than obtained with the standard Bonferroni correction or BH-False Discovery Rate methodologies ^28^. To select the most relevant functional categories, the term size was adjusted to 0 as minimum and 300 as maximum, avoiding redundant and generic terms.

### Statistical analysis

GraphPad Prism software 9.4.1 was used for the statistical analysis. Appropriate tests were chosen according to the data distribution based on the Saphiro-Wilk normality test. Outliers were identified and excluded by the ROUT method.

Comparisons for measured parameters and repeated measurements (RM) are indicated in each figure legend. Differences were analyzed by unpaired Student *t*-test, Mann-Whitney test, and one-way ANOVA or two-way ANOVA tests, and multiple comparisons were analyzed with the Tukey or Šídák post hoc tests, as appropriate; the tests used are indicated in each figure legend. Differences were considered statistically significant at p<0.05.

The number of animals used is indicated in the corresponding figures or figure legends. No statistical method was used to predetermine sample size. Sample size was chosen empirically according to our experience in the calculation of experimental variability. All experiments were carried out with at least 3 biological replicates.

Experimental groups were balanced in terms of animal age and weight. Only male mice were used because the *Myh11-Cre^ERT2^* transgene was inserted in the Y chromosome (https://www.jax.org/strain/019079). Investigators were not blinded to group allocation during experiments or to outcome assessments. Animals were genotyped before experiments, caged together, and treated in the same way.

## RESULTS

### Smooth muscle Cn is required for AAA formation

To investigate the selective contribution of EC-expressed and VSMC-expressed Cn to pathological VWR, we generated tamoxifen-inducible knockout mice conditionally lacking Cn in ECs or smooth muscle cells (SMCs). The catalytic subunit CnA is encoded by 3 genes (*Ppp3ca, Ppp3cb and Ppp3cc*), and the regulatory subunit CnB by 2 genes (*Ppp3r1* and *Ppp3r2*). The CnB2 subunit is expressed only in testes, whereas CnB1 is ubiquitously expressed and is indispensable for maintaining Cn activity ^29^. CnA and CnB are obligate heterodimers, such that if CnB is missing, CnA becomes unstable and is degraded ^30^. We therefore crossed LoxP-flanked *Ppp3r1* (*Cnb1*^f/f^) mice ^13^ with mice expressing tamoxifen-inducible Cre recombinase (Cre^ERT2^) specifically in ECs (*Cdh5*-Cre^ERT2^) ^14^ or in SMCs (*Myh11*-Cre^ERT2^) ^15^, obtaining *Cnb1*^f/f^;*Cdh5*-Cre^ERT2^ mice (EC-Cn^f/f^) or *Cnb1*^f/f^;*Myh11*-Cre^ERT2^ mice (SM-Cn^f/f^), respectively. *Cnb1*^f/f^-Cre^ERT2-neg^ together with *Cnb1*^wt/wt^;*Cdh5*-Cre^ERT2^ or *Cnb1*^wt/wt^;*Myh11*-Cre^ERT2^, were used as controls (Cn-Ctl) for EC-KO or SM-KO, respectively. Tamoxifen treatment induced *Cnb1* deletion specifically in ECs (EC-Cn^-/-^ mice) or SMCs (SM-Cn^-/-^ mice), as confirmed by immunoblot analysis of CnB expression in mouse aortic ECs (mAECs) from EC-Cn^-/-^ mice (**Figure 1A**) or in aortic extracts from SM-Cn^-/-^ mice (**Figure 1B**). Destabilization of the catalytic subunit CnA in the absence of the regulatory subunit CnB has been demonstrated in *in vivo* models ^19,30^. As expected, *Cnb1* deletion caused a drop in CnA protein expression (**Figure 1A-B**). Of note, Cn deficiency in ECs or SMCs did not substantially affect blood pressure (BP), aortic diameter, heart rate, or aortic tissue organization at baseline (**Figure 1C-E**).

**Figure 1.**
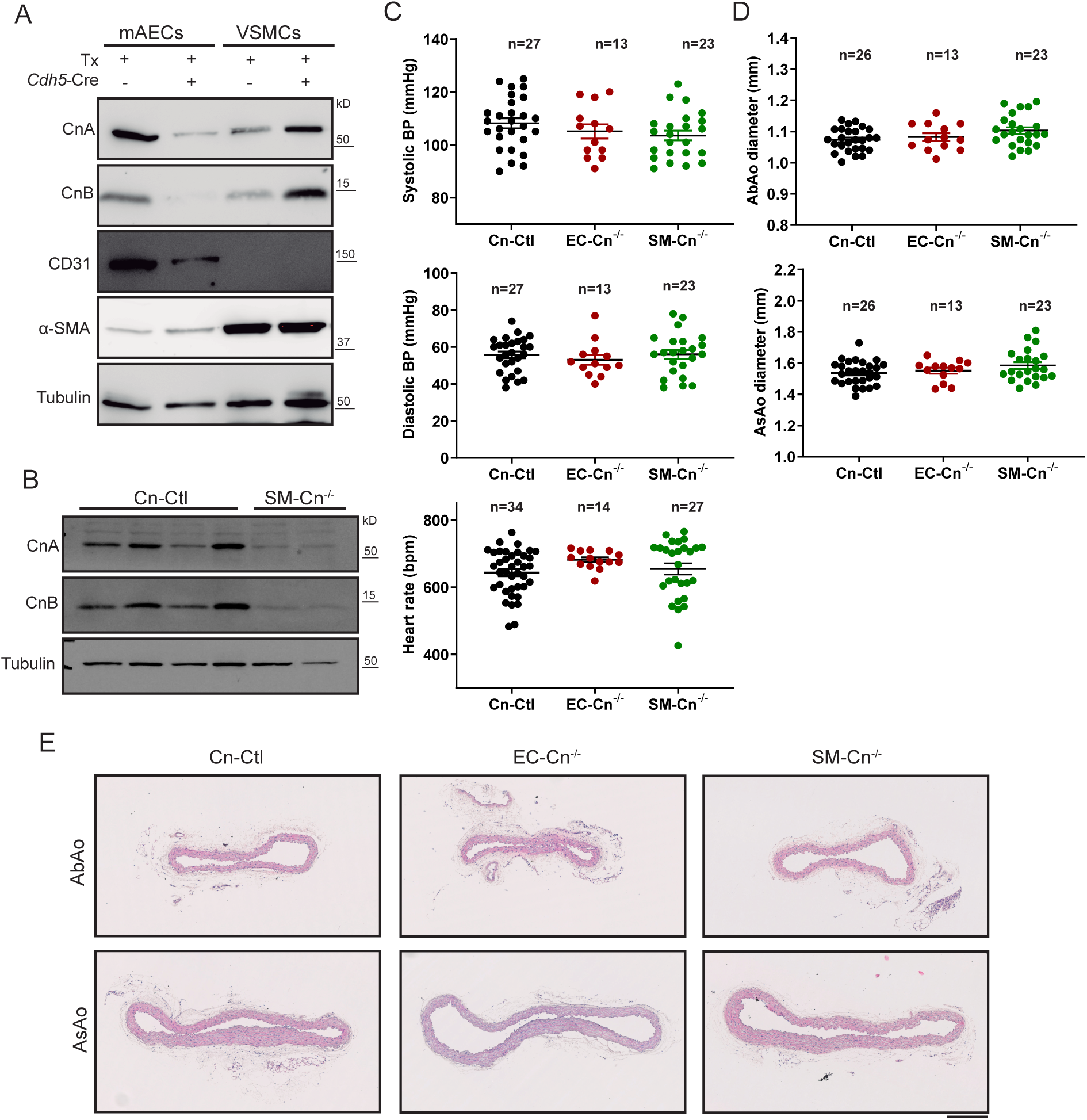
Conditional Cn deletion causes no evident symptoms in the absence of Ang-II. (**A**) Representative immunoblot analysis of CnA, CnB, CD31, α-SMA, and tubulin (loading control) in mouse aortic endothelial cells and vascular smooth muscle cells (n=4 independent experiments). (**B**) Representative immunoblot analysis of CnA, CnB, and tubulin (loading control) in total aortic extracts from 5 Cn-Ctl and 5 SM-Cn^-/-^ mice. (**C**) Systolic BP, diastolic BP, and heart rate (beats per minute, bpm), and (**D**) AbAo and AsAo diameters in Cn-Ctl, SM-Cn^-/-^, and EC-Cn^-/-^ mice. Each data point denotes an individual mouse, and the horizontal bars denote the mean (long bar) and s.e.m. The number of mice per group is indicated in each panel. (**E**) Representative images of hematoxylin-eosin–stained AbAo and AsAo cross sections from the indicated mice (n=5 mice per genotype). Scale bar, 250μm.

To assess the contributions of EC-expressed and SMC-expressed Cn to AAA, we used a mouse model ^16^ based on the infusion of Ang-II in mice previously fed a high-fat diet (HFD) for 12 weeks (**Figure 2A**). After 4 weeks of Ang-II treatment, a significant abdominal aorta (AbAo) dilation was observed in Cn-Ctl mice (**Figure 2B-D**). While Ang-II-induced AbAo dilatation was unaffected by deletion of Cn in ECs, it was abolished by Cn deletion in SMCs (**Figure 2C-D**). Cn deletion in ECs or SMCs did not prevent HFD-induced obesity in these mice (**<u>Suppl. Figure 1</u>**). Ex vivo images and histological analysis of AbAo cross sections revealed AbAo remodeling characterized by increased wall thickness, collagen deposition, and elastic fiber disarray in Ang-II-treated Cn-Ctl and EC-Cn^-/-^ mice that was not present in the AbAo of SM-Cn^-/-^ mice (**Figure 2E-G**). Together, these results indicate that SMC Cn, but not EC Cn, is critical for Ang-II-induced AbAo wall remodeling and AAA formation.

**Figure 2.**
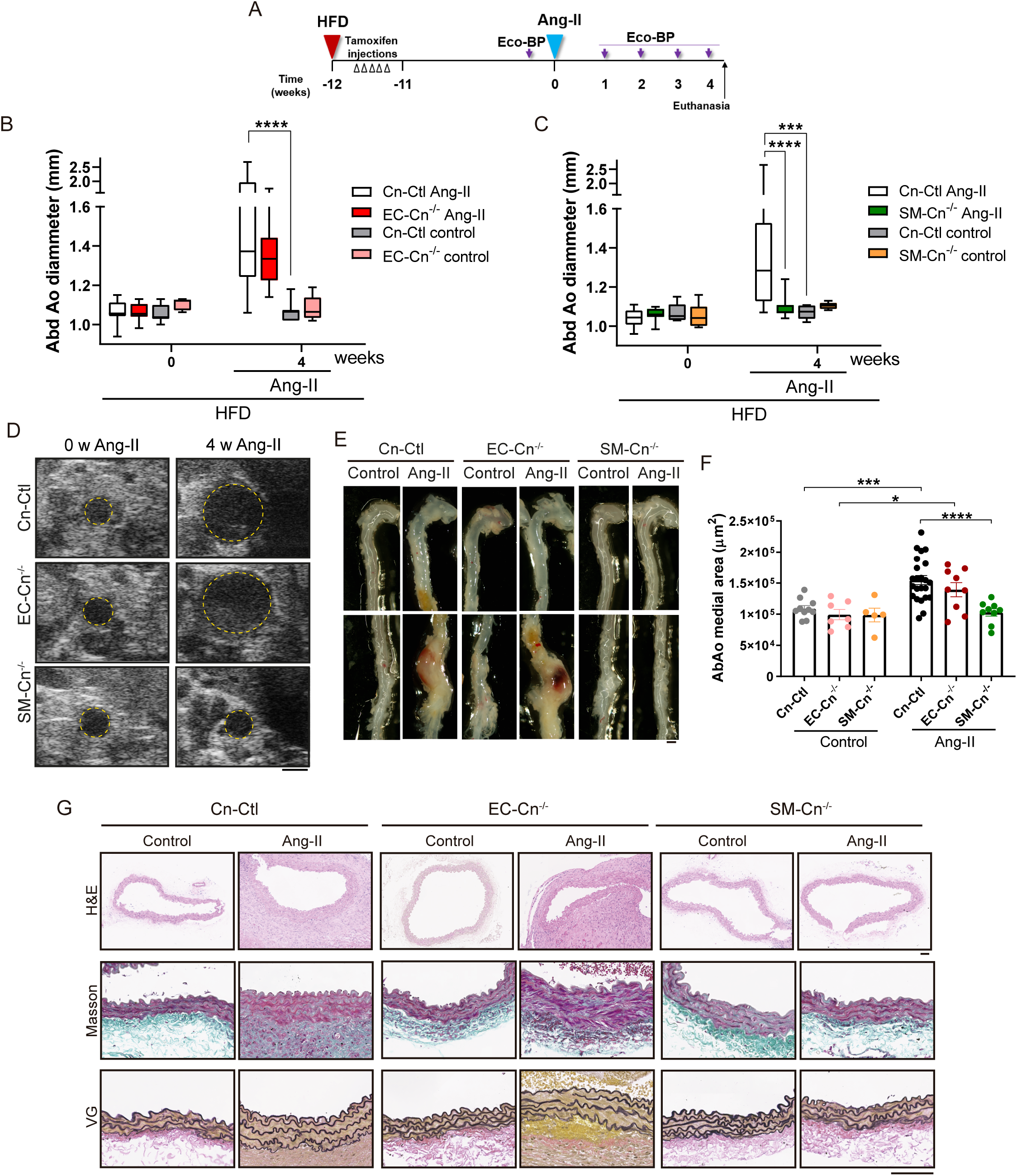
SMC Cn is required for Ang-II-induced AAA formation. (**A**) Experimental design: 10-12-week-old mice were treated with tamoxifen for 5 consecutive days (open arrowheads) during the first week of HFD feeding. After 12 weeks of HFD, Ang-II osmotic minipumps were implanted for 4 weeks; control mice were operated without minipump implantation. Maximum aortic diameter and BP were measured (Eco-BP) at the indicated time points (purple arrows), and mice were euthanized at the end of the experiment. (**B,C**) AbAo diameter in (**B**) Cn-Ctl (n=16), EC-Cn^-/-^ (n=12) Ang-II-treated and Cn-Ctl (n=7), EC-Cn^-/-^ (n=8) control mice, and in (**C**) Cn-Ctl (n=18), SM-Cn^-/-^ (n=16) Ang-II-treated and Cn-Ctl (n=8), SM-Cn^-/-^ (n=6) control mice. Data are presented as mean ± s.e.m. ****p<0.0001, ***p<0.001; repeated-measurements (RM) two-way ANOVA with Tukey’s post hoc test. (**D**) Representative ultrasound images of AbAo from mice before and after 4 weeks of Ang-II treatment. Yellow lines mark the lumen boundary. Scale bar, 1 mm. (**E**) Representative images of AAA. Scale bar, 1 mm. (**F**) Medial area in AbAo sections from Cn-Ctl (n=23), EC-Cn^-/-^ (n=9), and SM-Cn^-/-^ (n=9) Ang-II-treated mice and from Cn-Ctl (n=10), EC-Cn^-/-^ (n=7), and SM-Cn^-/-^ (n=5) control mice. Scale bar, 100µm. Each data point denotes an individual mouse, and data in histograms are presented as mean ± s.e.m. ****p<0.0001, ***p<0.001, *p<0.05; RM two-way ANOVA with Tukey’s post hoc test. (**G**) Representative images of hematoxylin-eosin (HE), Masson trichrome (Masson) and Van Giemson (VG) staining on AbAo sections from 4 Cn-Ctl, 3 EC-Cn^-/-^, and 4 SM-Cn^-/-^ mice. Scale bar, 100µm.

### SMC Cn is required for Ang-II-induced hypertension

Ang-II induces vasoconstriction and hypertension in mice, and we therefore investigated whether Cn deficiency affected Ang-II-induced hypertension in the same mice used for the experiments presented in Figure 2. While Cn-Ctl and EC-Cn^-/-^ mice showed a sharp and sustained increase in systolic and diastolic BP after 1 week of treatment with Ang-II, SM-Cn^-/-^ mice remained normotensive (**Figure 3**). These results strongly suggest that SMC Cn is a critical mediator of Ang-II-induced hypertension.

**Figure 3.**
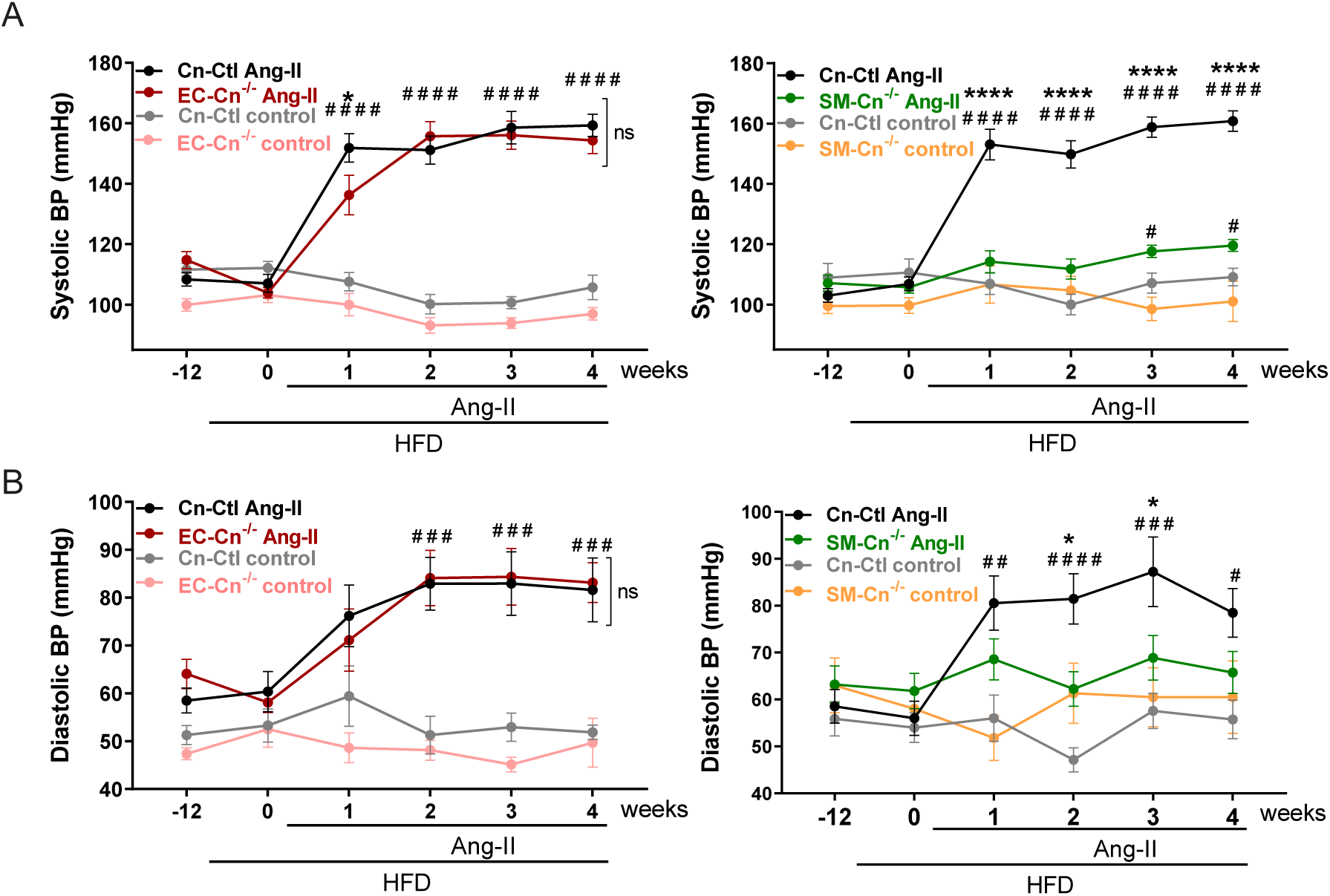
Cn deletion in SMCs prevents Ang-II-driven arterial hypertension in mice fed a HFD. (**A**) Systolic and (**B**) diastolic BP values at the indicated times in the same mice shown in Figure 2. (left) Cn-Ctl (n=16), EC-Cn^-/-^ (n=12) Ang-II-treated and Cn-Ctl (n=7), EC-Cn^-/-^ (n=8) control mice, and (right) Cn-Ctl (n=18), SM-Cn^-/-^ (n=16) Ang-II-treated and Cn-Ctl (n=8), SM-Cn^-/-^ (n=6) control. Data are presented as mean ± s.e.m. ****p<0.0001, *p<0.05 vs SM-Cn^-/-^ Ang-II or vs EC-Cn^-/-^ Ang-II, ^####^p<0.0001, ^###^p<0.001, ^##^p<0.01 and ^#^p<0.05 vs Cn-Ctl control, and (A, right) ^#^p<0.05 vs SM-Cn^-/-^ control; RM two-way ANOVA with Tukey’s post hoc test.

In rabbits, a high-cholesterol diet alters Ang-II signaling in the aorta ^31^. Therefore, to exclude possible secondary effects of HFD feeding on BP regulation by SMC Cn, we investigated the contribution of vascular Cn to Ang-II-induced hypertension in mice fed a chow diet. Infusion of Ang-II in tamoxifen-treated mice (**Figure 4A**) sharply increased BP in Cn-Ctl and EC-Cn^-/-^ mice from the first week of treatment but failed to substantially raise BP in SM-Cn^-/-^ mice (**Figure 4B <u>and Suppl. Figure 2</u>**). Similarly, Ang-II induced AbAo dilatation in Cn-Ctl and EC-Cn^-/-^ mice from the first week of treatment but did not have this effect in SM-Cn^-/-^ mice (**Figure 4C**). These results support a key role for SMC Cn in Ang-II-induced hypertension and AbAo dilatation.

**Figure 4.**
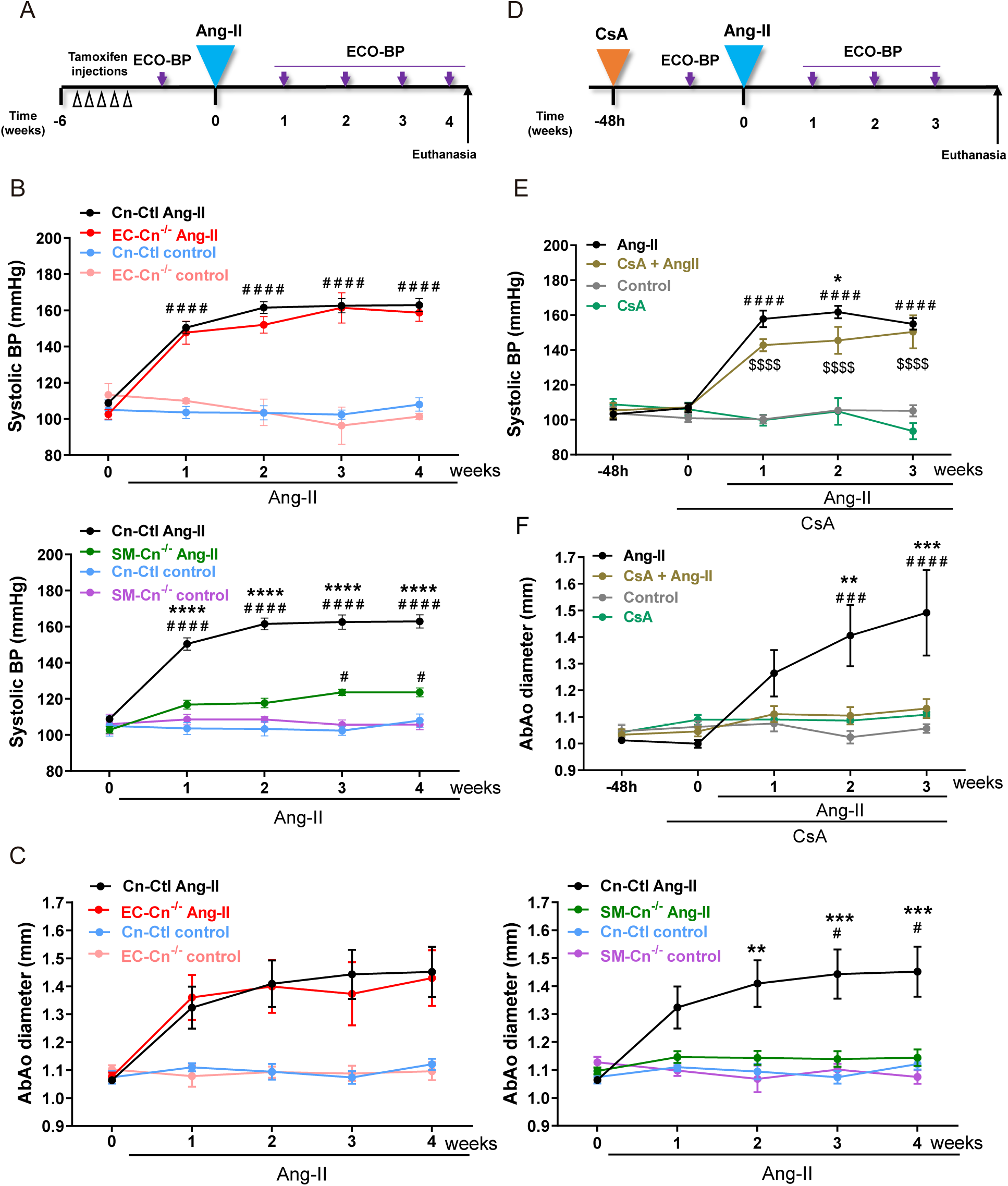
SMC Cn mediates Ang-II-induced hypertension independently of its phosphatase activity. (**A**) Experimental design: 10-12-week-old mice were treated with tamoxifen for 5 consecutive days (open arrow heads) and, after 6 weeks, Ang-II osmotic minipumps were implanted for 4 weeks in one group of mice; control mice were operated without minipump implantation. Maximum aortic diameter and BP were measured (Eco-BP) at the indicated time points (purple arrows), and mice were euthanized at the end of the experiment. (**B**) Systolic BP and (**C**) maximal AbAo diameters at the indicated times in Cn-Ctl (n=22), EC-Cn^-/-^ (n=10), and SM-Cn^-/-^ (n=18) Ang-II-treated mice and in Cn-Ctl (n=5), EC-Cn^-/-^ (n=3), and SM-Cn^-/-^ (n=5) control mice. Data are presented as mean ± s.e.m. ****p<0.0001, ***p<0.001, **p<0.01 vs SM-Cn^-/-^ Ang-II, ^####^p<0.0001 vs Cn-Ctl control, ^#^p<0.05 vs SM-Cn^-/-^ or Cn-Ctl control; RM two-way ANOVA with Tukey’s post hoc test. (**D**) Experimental design: 10-12-week-old mice were fitted with CsA osmotic minipumps; control mice were operated without minipump implantation. After 48h, Ang-II minipumps were implanted in a group of CsA-treated mice (CsA+Ang-II) and in a group of CsA control mice (Ang-II). The remaining CsA control mice were operated without minipump implantation (Control). Maximal aortic diameter and BP were measured (Eco-BP) at the indicated time points (purple arrows), and mice were euthanized at the end of the experiment. (**E**) Systolic BP and (**F**) maximal AbAo diameters at the indicated times in Ang-II-(n=10), CsA+Ang-II-(n=8), Control-(n=8), and CsA-(n=4) treated mice. Data are presented as mean ± s.e.m. ***p<0.001, **p<0.01, *p<0.05 vs CsA+Ang-II, ^####^p<0.0001, ^###^p<0.001 vs control, ^$$$$^p<0.0001 vs CsA; RM two-way ANOVA with Tukey’s post hoc test.

To investigate whether Cn deletion in SMCs also conferred resistance to hypertension induced by other factors, we treated mice with norepinephrine (NE) or the nitric oxide synthase inhibitor L-NAME (**<u>Suppl. Figure 3A</u>**). NE and L-NAME both markedly increased BP in Cn-Ctl mice relative to similarly treated SM-Cn^-/-^ mice and untreated controls (**<u>Suppl. Figure 3B</u>**). These results indicate that SMC-expressed Cn is an essential regulator of BP in response to diverse hypertensive stimuli.

#### Ang-II-induced hypertension does not require Cn phosphatase activity

A major role for Cn in BP regulation was unexpected because inhibition of Cn activity does not prevent Ang-II-induced hypertension in mice ^32^ and because long-term treatment of patients with Cn inhibitors causes hypertension, likely as a consequence of nephrotoxicity ^33^. A number of factors could account for this discrepancy, including different contributions of Cn to BP regulation in humans and mice, off-target effects of Cn inhibitors, or a distinct effect of pharmacological inhibition versus protein deletion. Pretreatment of mice with the Cn inhibitor CsA did not protect against Ang-II-induced hypertension in WT mice (**Figure 4D-E**); in contrast, CsA pretreatment did block Ang-II-induced AbAo dilatation in these mice (**Figure 4F**). This result confirms effective phosphatase inhibition and is consistent with a previous report showing that CsA prevents AbAo dilatation and AAA ^7^. Demonstrating efficient systemic inhibition of Cn phosphatase activity by CsA, NFATc3, a known Cn substrate, was hyperphosphorylated in the thymus of CsA-treated mice (**<u>Suppl. Figure 4A</u>**). Also consistent with previous reports ^32^, CsA pretreatment significantly reduced Ang-II-induced cardiac hypertrophy in these mice (**<u>Suppl. Figure 4 B-C</u>**). Together, these data indicate that CsA does not prevent Ang-II-induced hypertension and thus suggest that the contribution of Cn to Ang-II-induced hypertension is unrelated to its phosphatase activity.

To confirm the non-involvement of Cn phosphatase activity in Ang-II-induced hypertension, we inhibited Cn activity *in vivo* by inoculating WT mice with a lentivirus (LV) encoding the LxVP peptide fused to GFP (LV-LxVP). Like CsA, the LxVP peptide blocks the binding of Cn to its substrates, inhibiting its activity ^34^. Six weeks after LV inoculation, Ang-II was administered to mice inoculated with LV-LxVP or with LV encoding an inactive mutant version (LV-LxVPmut), and BP was determined (**Figure 5A**). Ang-II markedly increased systolic and diastolic BP in mice transduced with LV-LxVPmut or LV-LxVP, with no substantial difference between treatment groups (**Figure 5B**). Efficient transduction of all aortic layers was confirmed by GFP immunostaining of aortic sections (**Figure 5C**). To assess whether LV-LxVP efficiently blocked Cn/NFAT activation by Ang-II in these mice, we analyzed the subcellular localization and DNA-binding activity of NFAT proteins. NFATs are dephosphorylated by Cn and translocate to the nucleus in active form to bind target gene promoters. *In situ* southwestern histochemistry detected abundant active NFAT in cell nuclei in medial AbAo and AsAo sections from Ang-II-treated mice expressing LV-LxVPmut, but a nuclear NFAT signal was barely detected in sections from LV-LxVP transduced animals (**Figures 5D-E**). These data further support the idea that SMC Cn mediates Ang-II-induced hypertension independently of its phosphatase activity.

**Figure 5.**
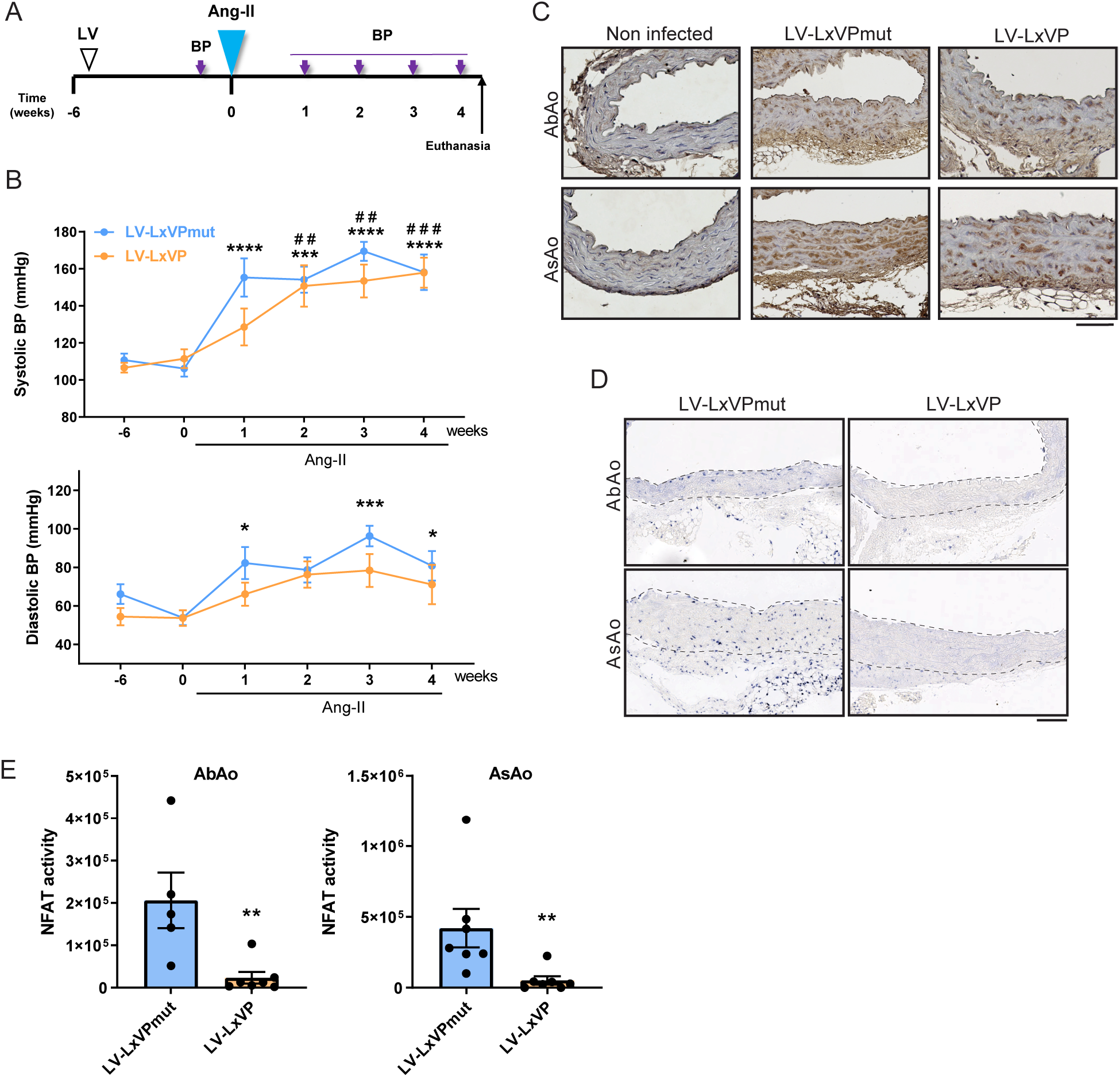
Cn inhibition by LxVP lentiviral transduction impairs NFAT activation *in vivo* without preventing Ang-II-induced hypertension. **(A)** Experimental design: 10-12-week-old mice were inoculated with LxVPmut-or LxVP-encoding lentivirus (LV) and, after 6 weeks, Ang-II osmotic minipumps were implanted for 4 weeks. BP was measured at the indicated time points (purple arrows), and mice were euthanized at the end of the experiment. (**B**) Systolic and diastolic BP in mice transduced with LV-LxVPmut (n=7) and LV-LxVP (n=7), shown as mean ± s.e.m. ****p<0.0001, ***p<0.001, **p<0.01, *p<0.05 and ^###^p<0.001, ^##^p<0.01 vs beginning of treatment (0 weeks) for LV-LxVPmut-and LV-LxVP-transduced mice, respectively; RM two-way ANOVA with Tukey’s post hoc test. (**C**) Representative images of GFP immunostaining in AbAo and AsAo cross sections from the indicated mice. (**D**) Representative images of southwestern histochemistry with an NFAT probe of AbAo and AsAo cross sections from the indicated mice and (**E**) quantification of the relative staining (NFAT activity) in the AbAo (left panel) and the AsAo (right panel). Dashed lines indicate the medial layer. Only mice with positive aortic GFP staining and consistent NFAT activity were included. Each data point denotes an individual mouse, and data in histograms are presented as mean ± s.e.m. Mann-Whitney test **p<0.01. Scale bars, 100 μm.

### SMC Cn is required for both early onset and long-term sustainability of Ang-II-induced hypertension

To investigate whether smooth muscle Cn was required only for the long-term maintenance of Ang-II-induced hypertension or also for its onset, we measured BP in Cn-Ctl and SM-Cn^-/-^ mice early after Ang-II infusion (**Figure 6A**). A robust increase in BP was observed in Cn-Ctl mice as early as 2h after Ang-II infusion and was maintained after 24h, whereas SM-Cn^-/-^ mice remained normotensive (**Figure 6B <u>and Suppl. Figure 5A</u>**). In parallel studies, WT mice were treated with CsA for 5 days before Ang-II infusion (**Figure 6C**). Consistent with the results of the long-term experiment with CsA (**Figures 4D-E**), CsA pre-treatment failed to prevent Ang-II-induced hypertension after 2h or 24h (**Figure 6D <u>and Suppl. Figure 5B</u>**). Efficient inhibition of Cn phosphatase activity in CsA-treated mice was confirmed by the NFAT phosphorylation status in the thymus and the inhibition of aortic *Rcan1-4* mRNA expression (**<u>Suppl. Figure 5C-D</u>**). These results strongly suggest that Ang-II activates molecular mechanisms critical for the early increase in BP that require Cn expression in SMCs, but not its phosphatase activity.

To determine whether the presence of Cn is required for long-term, sustained hypertension, we triggered Cn deletion in SMCs 7 days after inducing hypertension with Ang-II (**Figure 6E**). Treatment with Ang-II for 7 days elicited a similar degree of hypertension in Cn-Ctl and SM-Cn^f/f^ mice (**Figure 6F**). At this point, all mice were treated with tamoxifen (Tx) to elicit smooth muscle deletion of Cn in the SM-Cn^f/f^ mice, generating SM-Cn^-/-^ mice. BP decreased in SM-Cn^-/-^ mice as early as 1 day after the first Tx administration and continued to decline until the end of the experiment; in contrast, Cn-Ctl mice remained hypertensive until the end of the experiment (**Figure 6F**). These data strongly suggest that smooth muscle Cn is actively involved in the onset and maintenance of Ang-II-induced hypertension, identifying smooth muscle Cn as a potential target for the treatment of hypertension.

**Figure 6.**
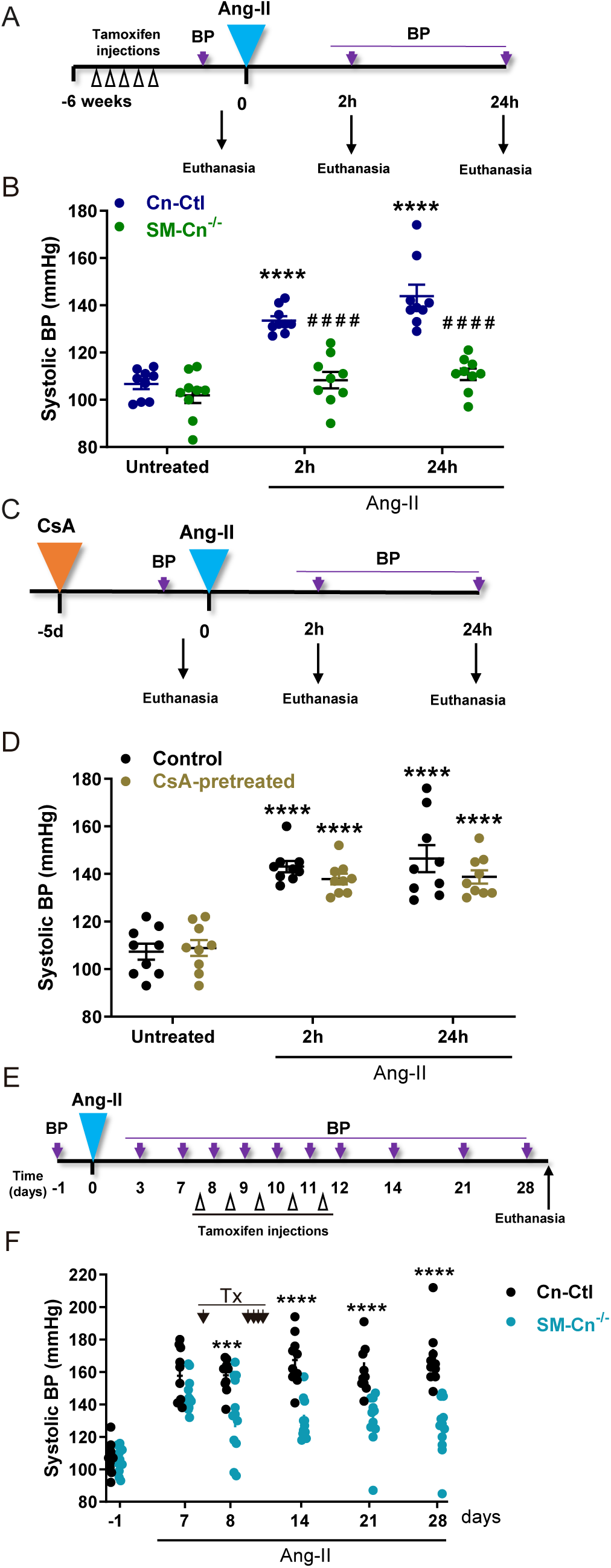
SMC Cn is required for both the onset and the long-term sustainability of Ang-II-induced hypertension. (**A**) Experimental design: 10-12-week-old mice were treated with tamoxifen for 5 consecutive days (open arrow heads) and, after 6 weeks, Ang-II osmotic minipumps were implanted for 2h and 24h. BP was measured at the indicated time points (purple arrows), and mice were euthanized and analyzed when indicated. (**B**) Systolic BP values in the indicated mice (n=9 mice per group and time point). Each data point denotes an individual mouse, and the horizontal bars denote the mean (long bar) and s.e.m. ****p<0.0001 vs baseline; RM two-way ANOVA with Tukey post hoc test. ^####^p<0.0001 vs 2h or 24h Ang-II Cn-Ctl; RM two-way ANOVA with the Šídák post hoc test. (**C**) Experimental design: 10-12-week-old mice were fitted with CsA osmotic minipumps (CsA-pretreated); control mice were operated without minipump implantation (Control). After 5 days, Ang-II minipumps were implanted in CsA-pretreated and Control mice for 2h and 24h. BP was measured at the indicated time points (purple arrows), and mice were euthanized and analyzed when indicated. (**D**) Systolic BP in the indicated mice (n=9 mice per group and time point). Each data point denotes an individual mouse, and the horizontal bars denote the mean (long bar) and s.e.m. ****p<0.0001 vs baseline; RM two-way ANOVA with Tukey’s post hoc test. (**E**) Experimental design: 10-12-week-old mice were fitted with Ang-II osmotic minipumps for 4 weeks. After 7 days of Ang-II infusion, tamoxifen was administered for 5 consecutive days (open arrow heads). BP was measured at the indicated time points (purple arrows), and mice were euthanized at the end of the experiment. (**F**) Systolic BP measurements at the indicated times in Cn-Ctl (n=10) and SM-Cn^-/-^ (n=12) mice. Each data point denotes an individual mouse, and the horizontal bars denote the mean (long bar) and s.e.m. ****p<0.0001, ***p<0.001 vs Cn-Ctl Ang-II; RM two-way ANOVA with Šídák post hoc test. Tx indicates tamoxifen administration time points.

### SMC Cn orchestrates the Ang-II-induced transcriptional program involved in arterial contractility and hypertension

To gain insight into the Cn-mediated molecular mechanisms underlying the early onset of hypertension, we performed a transcriptomic analysis to identify genes whose regulation by Ang-II in the aorta requires Cn expression in SMCs. A heatmap representation of normalized average expression shows that smooth muscle Cn is required for the differential expression of most of the genes regulated by Ang-II in the aorta (**Figure 7A**). Our analysis identified 800 differentially expressed genes (DEGs) in the aortas of Cn-Ctl mice treated for 2h with Ang-II versus untreated mice; of these DEGs, almost 90% (722 genes) showed no evidence of Ang-II regulation in the aortas of SM-Cn^-/-^ mice (**<u>Suppl. Figure 6A</u>**). To narrow down the number of candidate mediators of Ang-II-induced hypertension, we focused on a group of 336 DEGs that were also differentially expressed between Ang-II-treated Cn-Ctl and SM-Cn^-/-^ mice (**<u>Suppl. Figure 6B</u>**).

**Figure 7.**
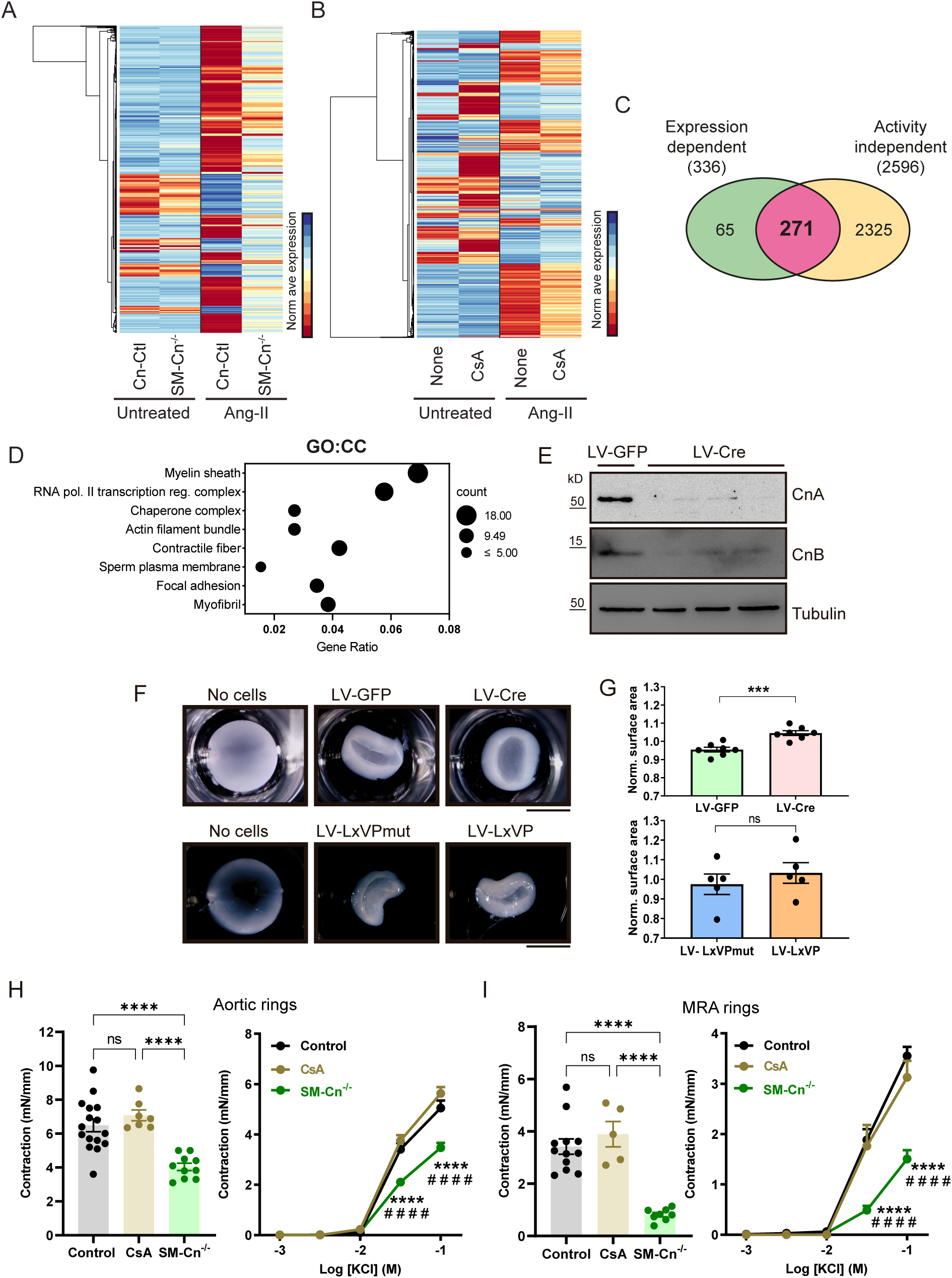
SMC Cn orchestrates an Ang-II-induced transcriptional program related to arterial contractility. (**A,B**) Heatmaps of the normalized average expression of DEGs regulated by Ang-II relative to the untreated conditions in each RNAseq dataset. (**C**) Venn diagram comparing the number of Cn expression-dependent DEGs (336) and Cn activity-independent DEGs (2596), and showing the number of shared DEGs whose regulation by Ang-II requires Cn expression but not its activity (271). (**D**) Most significantly enriched cellular components (CC) of the 271 Cn expression-dependent DEGs, obtained by g:Profiler. (**E**) Representative immunoblot analysis of CnA, CnB, and tubulin (loading control) in protein extracts from LV-GFP-and LV-Cre-transduced primary VSMCs (n=7 individual experiments). Molecular weights (kDa) are indicated. (**F**) Representative images and (**G**) normalized surface area of fixed collagen gels under the indicated conditions. Each data point denotes the mean of each experiment, and data in histograms are presented as mean ± s.e.m. (n=7 independent experiments with LV-GFP and LV-Cre; n=5 independent experiments with LV-LxVP and LV-LxVPmut). ***p<0.001; n.s., non-significant; unpaired Student t-test. Scale bars, 5 mm. Quantification of tension in (**H**) aortic rings and (**I**) MRA rings from control (Cn-Ctl or WT), SM-Cn^-/-^, or CsA-treated WT mice after stimulation for 30 minutes with 80 mM KCl (left panels) and the indicated KCl concentrations (right panels). Left panels, each data point denotes an individual mouse, and data in histograms are presented as mean ± s.e.m. ****p<0.0001; one-way ANOVA with Tukey post hoc test. Right panels, data are shown as mean ± s.e.m. ****p<0.0001 vs control, ^####^p<0.0001 vs CsA; two-way ANOVA with Tukey post hoc test. Aortic rings from 16 control, 10 SM-Cn^-/-^, and 7 CsA-treated mice and MRA rings from 12 control, 8 SM-Cn^-/-^, and 5 CsA-treated mice were used.

Since CsA pretreatment did not prevent Ang-II-induced hypertension, we reasoned that genes whose induction by Ang-II was sensitive to CsA would be unlikely to mediate hypertension. We therefore performed an additional transcriptomic analysis to identify genes whose regulation by Ang-II was unaffected by CsA. The performance of this study was very high and identified 2653 DEGs in the aortas of mice treated with Ang-II for 2h (**<u>Suppl. Figure 6C</u>**). The heatmap representation of these DEGs showed a marked effect of Ang-II treatment on the aortic transcriptome and a subtle effect of CsA pretreatment on gene expression regulation by Ang-II (**Figure 7B**), with 2596 of the Ang-II-regulated genes (almost 98%) insensitive to CsA pretreatment (**<u>Suppl. Figure 6C</u>**). Comparison of these genes with the 336 Cn-dependent DEGs identified a group of 271 genes whose regulation by Ang-II required Cn expression but not its phosphatase activity (**Figure 7C**). Functional annotation clustering of these genes revealed enrichment in cluster terms strongly associated with cellular components involved in cell contractility regulation, such as “actin filament bundle”, “contractile fiber”, “focal adhesion”, and “myofibril” (**Figure 7D**).

VSMC contractility has a direct effect on BP regulation ^3^, and actomyosin cytoskeleton dynamics is critical for VSMC contractility regulation ^35^. To investigate the influence of Cn on VSMC contractile capacity, we cultured primary Cn^f/f^ VSMCs transduced with Cre-encoding or GFP-encoding LV on collagen disks. In this assay, contractile capacity is determined by measuring the disk surface area (**<u>Suppl. Figure 7</u>**). Transduction with Cre-encoding LV efficiently deleted Cn expression (**Figure 7E**) and substantially impaired gel contraction relative to cells transduced with LV-GFP (**Figures 7F-7G**). In contrast, transduction of WT VSMCs with LxVP-encoding LV failed to prevent gel contraction (**Figures 7F-7G**). These results suggest that Cn regulates VSMC contractility through a mechanism unrelated to its phosphatase activity. To investigate the contribution of SMC Cn to vessel contractility regulation in a more physiological setting, we measured the KCl-induced contraction of vehicle-or CsA-pretreated aortic and mesenteric resistance artery (MRA) rings from WT, Cn-Ctl, and SM-Cn^-/-^ mice. Cn deficiency, but not inhibition of its phosphatase activity with CsA, decreased aortic and MRA contractility in response to saturating (**Figures 7H-7I, <u>left panels</u>**) or increasing KCl concentrations (**Figures 7H-7I, <u>right panels</u>**), providing further support for the idea that Cn expression in SMCs is critical for arterial contractility regulation independently of Cn phosphatase activity.

To investigate the potential association of the group of 271 genes with hypertension, we performed an analysis using the Harmonizome dataset collection (http://amp.pharm.mssm.edu/Harmonizome). This analysis provided a hypertension score (HS) for >70% of these genes, suggesting a likely association with hypertension and supporting the validity of our transcriptomics approach as a way to identify genes encoding mediators of Ang-II-induced hypertension. Graphical representation of the fold change (FC) in expression induced by Ang-II for the genes with the highest HS revealed a markedly different regulation in SM-Cn^-/-^ mice relative to Cn-Ctl, WT, and CsA-pretreated mice (**Figure 8A**). We analyzed the expression of 3 of the genes with the highest HS in aortas from individual mice by RT-qPCR. The expression of *Serpine1, Ier3,* and *Gja1* after 2h exposure to Ang-II was markedly inhibited in Cn-deficient mice but was not repressed by treatment with CsA (**Figure 8B**), identifying these genes as candidate mediators of Ang-II-induced hypertension.

**Figure 8.**
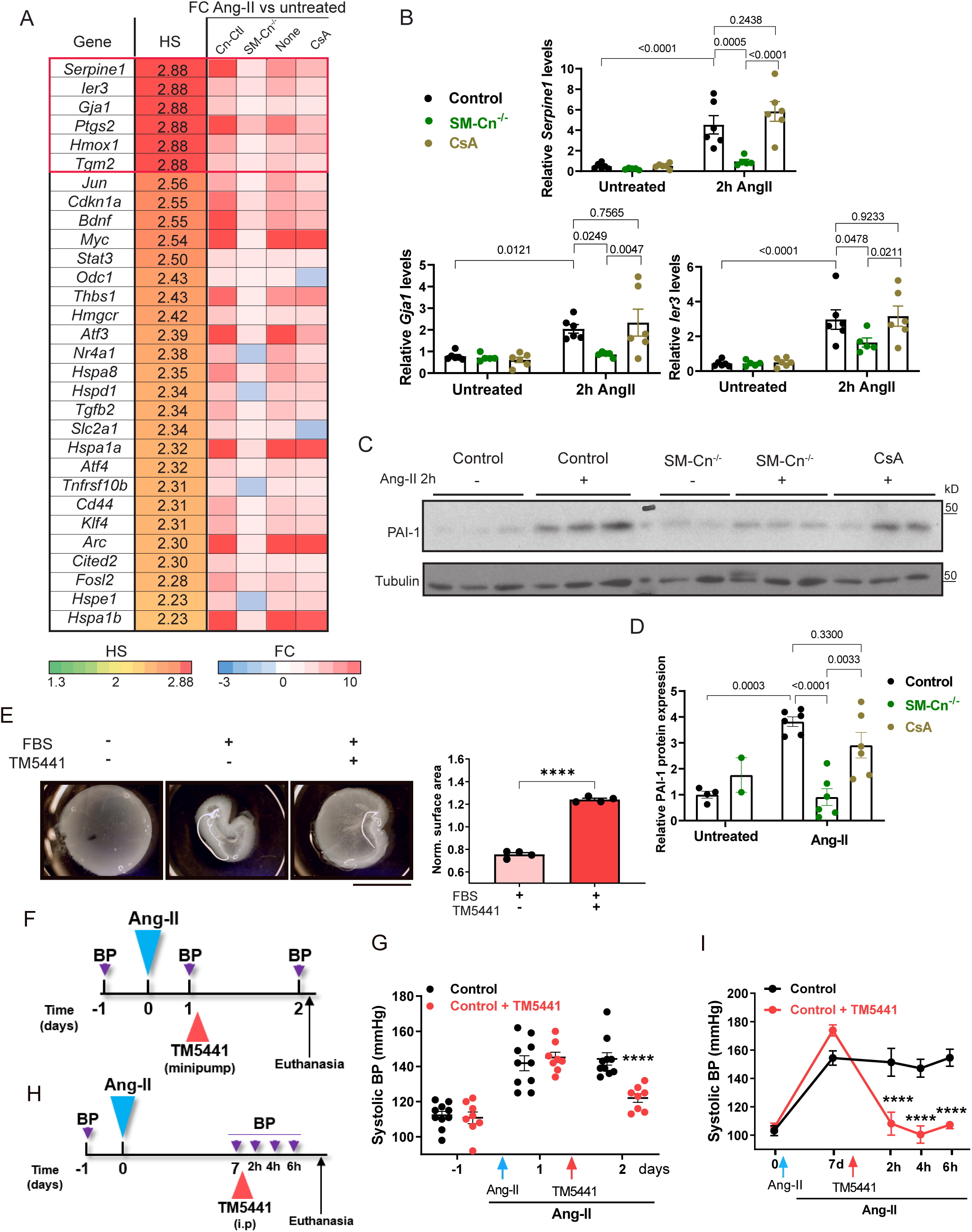
PAI-1 mediates VSMC contractility and Ang-II-induced hypertension. (**A**) List of the 30 genes with the highest hypertension score (HS), showing their gene symbol, HS, and expression fold change (FC) after 2h of Ang-II treatment relative to untreated mice. (**B**) RT-qPCR analysis of mRNA expression in aortic extracts from untreated control (3 Cn-Ctl plus 3 WT), SM-Cn^-/-^ (n=5), and CsA-pretreated (n=6) mice and from Ang-II-treated control (3 Cn-Ctl plus 3 WT), SM-Cn^-/-^ (n=5), and CsA-pretreated (n=6) mice. Two-way ANOVA with Šídák post hoc test; p values and comparisons are indicated. (**C**) Representative immunoblot analysis of PAI-1 and tubulin (loading control) and (**D**) quantification of their relative expression in protein extracts from untreated control (n=4) and SM-Cn^-/-^ (n=2) mice and from Ang-II-treated control (n=6), SM-Cn^-/-^ (n=6), and CsA-pretreated (n=6) mice. Molecular weights (kDa) are indicated. Each data point denotes an individual mouse, and data in histograms are presented as mean ± s.e.m. One-way ANOVA with Tukey post hoc test; p values and comparisons are indicated. (**E**) Representative images (left panels) and normalized surface area of fixed collagen gels (right panel) under the indicated conditions. Each data point denotes the mean of each experiment, and data in histograms are presented as mean ± s.e.m. (n=4 independent experiments with with 3% FBS and 50 µM TM5441). ****p<0,0001, unpaired Student t-test. Scale bars, 5 mm. (**F**) Experimental design: 10-12-week-old mice were fitted with Ang-II osmotic minipumps (Ang-II) 24h before implantating TM5441 minipumps in a group of Ang-II-pretreated mice (Ang-II+TM5441). BP was measured at the indicated time points (purple arrows), and mice were euthanized at the end of the experiment. (**G**) Systolic BP at the indicated times in 10 Ang-II-treated mice (Ang-II) and 8 mice treated with Ang-II+TM5441. Data are presented as mean ± s.e.m. ****p<0.001, vs Control RM two-way ANOVA with Šídák post hoc test. (**H**) Experimental design: 10-12-week-old mice were fitted with Ang-II osmotic minipumps (Ang-II) 7 days before intraperitoneally injecting TM5441 or vehicle to a group of Ang-II-pretreated mice. BP was measured at the indicated time points (purple arrows), and mice were euthanized at the end of the experiment. (**I**) Systolic BP at the indicated times in 5 Ang-II + vehicle (Control) and 6 mice treated with Ang-II+TM5441 (Control + TM5441). Data are presented as mean ± s.e.m. ****p<0.001, vs Control RM two-way ANOVA with Šídák post hoc test.

Consistent with the mRNA data, the expression of PAI-1, the protein encoded by *Serpine1*, was increased in aortas of mice treated with Ang-II for 2h and this increase did not occur in SM-Cn^-/-^ mice and was not prevented by the treatment with CsA (**Figure 8C-8D**). Given the identification of *Serpine1* as a major candidate mediator of Ang-II-induced hypertension, we investigated the potential contribution of PAI-1 to contractility regulation and Ang-II-induced hypertension. Pre-treatment of VSMCs with the PAI-1 inhibitor TM-5441 nearly abolished their capacity to contract collagen gels (**Figure 8E**). Furthermore, when mice were treated with this inhibitor 24 hours after the induction of hypertension with Ang-II (**Figure 8F-8G**), there was a marked reduction in BP 24 h after the inhibitor administration (**Figure 8G**). To validate the impact of PAI-1 inhibition *in vivo*, we employed an alternative experimental approach and found that the hypertension induced by 7 days of Ang-II treatment was completely reversed by the acute intraperitoneal administration of TM5441 (**Figure 8H-I**). These results further emphasize the association between BP regulation and SMC contractility and demonstrate that PAI-1, a protein whose expression regulation by Ang-II relies on Cn expression but does not necessitate its phosphatase activity, is a critical mediator of SMC contractility and Ang-II-induced hypertension.

## DISCUSSION

The results of this study reveal that the phosphatase activity of SMC Cn is critical for Ang-II-induced abdominal aorta dilatation but that SMC Cn unexpectedly drives hypertension independently of its phosphatase activity. Through this non-enzymatic mechanism, SMC Cn is the master regulator of a gene expression program that leads to hypertension. Almost 90% of the genes regulated by the vasopressor Ang-II during hypertension onset require Cn expression in SMCs, but only a small fraction of these genes require Cn phosphatase activity for this regulation. Together, our findings suggest a critical structural role for Cn in BP and gene expression regulation.

Cn was identified in previous studies as a mediator of Ang-II-induced AAA and vascular remodeling ^7,12,36^, yet the aortic cell types responsible for this role remained unknown. Our findings demonstrate that Ang-II-induced AAA formation in hyperlipidemic mice requires Cn expression in SMCs, but not in ECs. Our previous research indicated that AAA formation requires the phosphatase activity of Cn ^7^. With our new findings, we now show that, while the development of hypertension also requires the presence of Cn in SMCs, it does not require Cn enzymatic activity.

Hypertension is a complex and multifactorial disease and, despite its high prevalence, the basis of its pathogenesis is not fully understood, creating problems in its clinical management and therapy ^2^. The involvement of VSMCs in hypertension has been demonstrated in several studies ^37–40^. Our present results show that specific deletion of Cn in SMCs, but not in ECs, prevents Ang-II-induced increases in systolic and diastolic BP. Moreover, Cn deletion in SMCs also inhibited hypertension induced by the vasopressors NE and L-NAME. Since SMC Cn deletion 1 week after inducing hypertension with Ang-II substantially decreased BP, our results point to an essential role for SMC Cn not only in the onset but also in the maintenance of hypertension. A recent report suggested that the catalytic subunit CnAβ is required for sustained Ang-II-induced hypertension but not for its induction, since constitutive CnAβ deletion did not prevent Ang-II-induced hypertension during the first 3 weeks of treatment but did cause a decrease in systolic BP 1 week later ^36^. Since *Cnb1* deletion in SMCs leads to destabilization of both Cn catalytic subunits expressed in these cells (CnAα and CnAý) and blocks Ang-II-induced hypertension, we propose that while CnAα mediates the induction of hypertension by Ang-II, CnAβ may be required for its maintenance.

Inhibition of Cn phosphatase activity with the immunosuppressor CsA prevented Ang-II-induced aortic dilatation but did not prevent Ang-II-induced hypertension. The discrepancy between the effects of Cn inhibition and genetic ablation could be due to the reported hypertensive effect of long-term CsA treatment ^41–43^ or to off-target effects of the drug ^44^. However, we detected no BP increase in mice treated with CsA alone. It should be noted that the earlier reports used higher CsA doses (20-30 mg/kg/d, vs 5 mg/kg/d in this study) and different administration routes ^41–43^. These results suggest that SMC Cn mediates Ang-II-induced hypertension through a mechanism independent of its phosphatase activity, an idea supported by our finding that LVs encoding the Cn inhibitory peptide LxVP also failed to impede Ang-II-induced hypertension despite efficiently blocking activation of the Cn substrate NFAT.

Catalysis-independent roles have already been described for some enzymes; for example, short-term presynaptic plasticity requires the presence of αCamKII but not its kinase activity ^45^. Although Cn function has broadly been attributed to its phosphatase activity, a phosphatase-independent role has been proposed in cardiac fibrosis ^32^. Moreover, Cn interaction with its substrate NFAT leads not only to its dephosphorylation-mediated activation but also to its retention in the nucleus by masking the nuclear export signal ^46^. Several Cn-interacting proteins have recently been identified that could potentially mediate a structural role of Cn in the distribution of postsynaptic density ^47^, and new Cn interactors and Cn-binding sites distinct from the canonical LxVP and PIxIT sites have been identified ^48^. Some of the identified Cn interactors can serve as Cn scaffolds or regulate Cn enzymatic activity or its subcellular distribution ^49^, and our results suggest that some Cn interactors might mediate BP regulation independently of Cn phosphatase activity.

Although BP is largely determined by the resistance arteries, large arteries also contribute by regulating pulse wave pressure during the cardiac cycle, and changes in aortic stiffness or compliance are strongly associated with hypertension in humans ^50^. There is increasing evidence that large artery damage has a deleterious impact on the microcirculation that affects hypertension; however, the underlying molecular mechanisms are poorly understood ^4^. Since we detected hypertension as early as 2 h after Ang-II infusion in WT and Cn-Ctl mice, we reasoned that a transcriptomic analysis of the aorta performed at this time could identify genes mediating hypertension onset. The regulation of nearly 90% of the genes affected by Ang-II in the aorta required Cn expression in SMCs, indicating that among the numerous signaling pathways activated by Ang-II, those mediated by Cn are critical, at least in the aorta, for early signaling events. Consistent with our previous results showing that just 11 of >1500 Ang-II-regulated genes in cultured VSMCs are sensitive to Cn inhibition by CsA ^7^, only 2% of the transcriptome regulated by Ang-II in the aorta required Cn phosphatase activity. These results further support a structural, phosphatase-activity-independent role for Cn in gene expression regulation.

To identify candidate mediators of hypertension, we focused on genes whose regulation by Ang-II was dependent on Cn expression but not on its phosphatase activity. Functional annotation clustering analysis of these genes revealed that Cn orchestrates the induction of a gene expression program enriched in cellular components strongly associated with VSMC contractility regulation. Moreover, Cn deletion in cultured VSMCs, but not inhibition of its phosphatase activity, impaired the contractile capacity of these cells. Accordingly, Cn deletion in SMCs, but not its phosphatase activity inhibition, diminished the contractile capacity of aortic and resistance arteries. These results are consistent with previous findings showing that impaired vessel contractility leads to diminished Ang-II-induced hypertension ^15,37,39,40,51^ and strongly suggest that SMC Cn regulates artery contraction and hypertension onset and maintenance in a phosphatase-activity-independent manner.

Cn deletion prevented not only Ang-II-induced hypertension, but also the BP increase induced by norepinephrine or L-NAME. It should be noted that these 3 stimuli increase [Ca^2+^]_i_, which is critical for VSMC contraction, and that Cn is implicated in the regulation of calcium dynamics ^52,53^. Thus, one interesting possibility is that Cn might regulate BP by modulating [Ca^2+^]_i_ and therefore SM contraction. In this regard, a gene with the highest hypertension score (HS) in the Harmonizome analysis was *Gja1*, which encodes connexin 43 (Cx43), a protein that regulates the Ca^2+^-dependent contraction of VSMCs ^54^.

Our results indicate that SMC Cn plays a key structural role in orchestrating the early induction of a gene expression program that includes 271 genes. While many of these genes are related to vessel contractility and hypertension onset and maintenance, nearly 30% of the genes whose regulation by Ang-II requires Cn expression but not its enzymatic activity have not been previously linked to hypertension and are thus potential novel targets for therapeutic intervention in hypertension.

The identification of Serpine1 as one of the genes with the highest HS whose regulation by Ang-II is mediated by the structural role of SMC Cn is noteworthy because this gene encodes plasminogen activator inhibitor type-1 (PAI-1), whose plasma levels correlate with clinical hypertension ^55^. *Serpine1* deletion reduces L-NAME-induced hypertension ^56^ and our present results also show that PAI-1 inhibition virtually abolished the contractile capacity of VSMCs and reversed the hypertension induced by Ang-II. Taken together, these results dentify PAI-1 as a calcineurin-regulated mediator of hypertension and urge the evaluation of PAI-1 inhibitors for the treatment of hypertension.

## Acknowledgments

We thank S. Bartlett for English language editing and the CNIC histology, genomics, bioinformatics, and advanced imaging units for excellent technical support and advice. The CNIC is supported by the Spanish Ministerio de Ciencia e Innovación and the Pro-CNIC Foundation; CBMSO is supported by Consejo Superior de Investigaciones Científicas and Universidad Autónoma de Madrid. CBMSO and CNIC are Severo Ochoa Centers of Excellence (CEX2021-001154-S and CEX2020-001041-S, respectively). The project leading to these results has received funding from “La Caixa” Banking Foundation under project codes HR18-00068 (to M.R.C. and J.M.R.) and HR17-00247 (to J.V.); grants PID2020-115217RB-I00 and PID2021-122388OB-I00 to M.R.C and J.M.R, respectively, funded by MCIN/AEI/10.13039/501100011033; the Instituto de Salud Carlos III (CIBER-CV CB16/11/00264); and Spanish Ministerio de Ciencia e Innovación fellowships FPI (BES-2016-076637) to P.S.Y.-L., FPI (BES-2016-077649) to S.M.-G., and Sara Borrell (CD18/00028) and Juan de la Cierva (IJC2020-044581-I) to M.T.

## Author Contributions

M.R.C. and J.M.R conceived the study; P.S.Y.-L., Y.S., S.M.-M., M.R.C., and J.M.R. designed the study and analyzed the data; P.S.Y.-L. and Y.S. performed most of the experiments, with contributions from S. M.-M., A.A., M.T., A.C., A.I.T., D.L.-M., and S.M.-G.; A.A. performed bioinformatics analysis under the supervision of J.V., who also provided ideas for the project; D.N.C. provided ideas for the project; P.S.Y.-L., J.M.R., and M.R.C. wrote the manuscript with contributions from S.M.-M and A.A. All authors read and approved the manuscript.

