## Supplemental Figures for "Phosphatase-independent activity of smooth-muscle calcineurin orchestrates a gene expression program leading to hypertension"

### SUPPLEMENTARY FIGURES

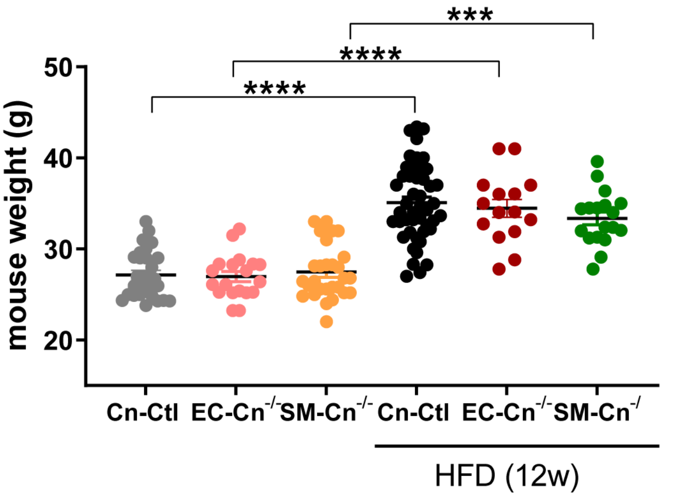


**Supplementary figure 1**. **Cn deletion in SMCs does not prevent HFD-induced body weight gain.** Body weight of 46 Cn-Ctl, 19 EC-Cn$^{-/-}$, and 18 SM-Cn$^{-/-}$ mice before and after 12 weeks of HFD. Each data point denotes an individual mouse, and the horizontal bars denote the mean (long bar) and the s.e.m. ****p<0.0001, ***p<0.001; one-way ANOVA with Tukey's multiple comparison post hoc test.

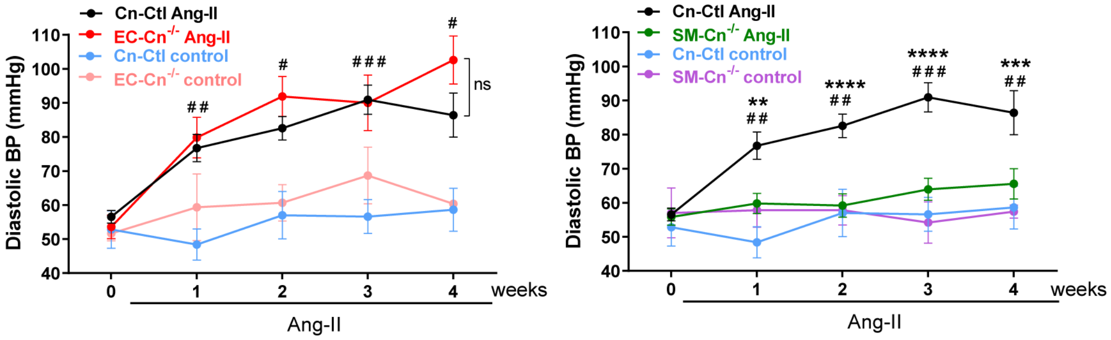


**Supplementary figure 2. Cn deletion in SMCs inhibits AngII-induced diastolic hypertension.** Diastolic BP values at the indicated times in Cn-Ctl (n=22), EC-Cn$^{-/-}$ (n=10), and SM-Cn$^{-/-}$ (n=18) Ang-II-treated mice and in Cn-Ctl (n=5), EC-Cn$^{-/-}$ (n=3), and SM-Cn$^{-/-}$ (n=5) control mice. Data are mean ± s.e.m. ****p<0.0001, ***p<0.001, **p<0.01 vs SM-Cn$^{-/-}$ Ang-II, $^{###}$p<0.001 vs Cn-Ctl control, $^{##}$p<0.01, $^{#}$p<0.05 vs SM-Cn$^{-/-}$ or Cn-Ctl control; RM two-way ANOVA with Tukey's post hoc test.

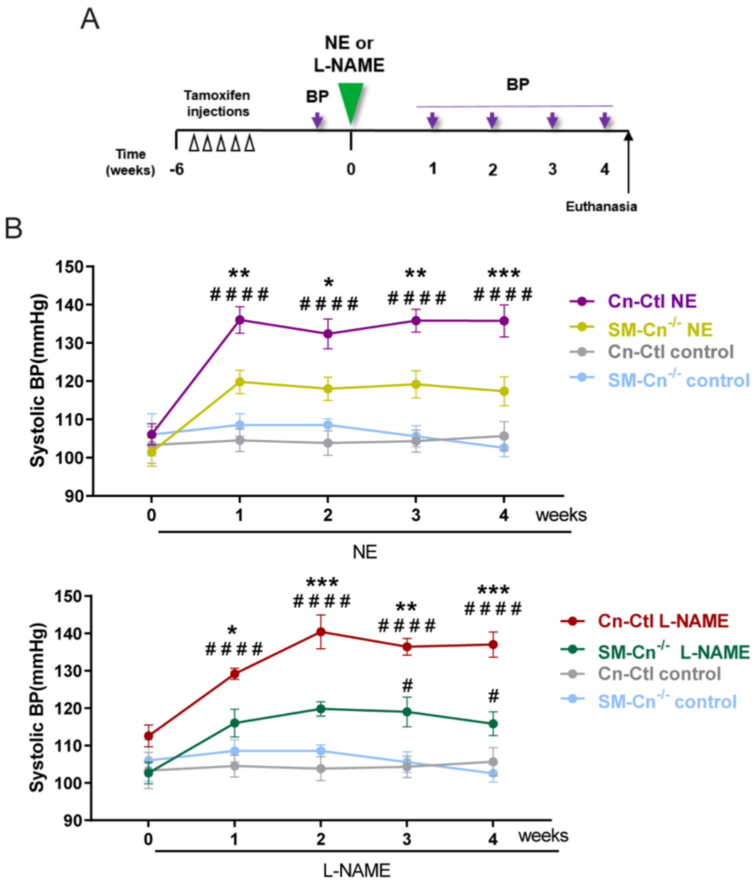


**Supplementary figure 3. Cn deletion in SMCs inhibits norepinephrine- and L-NAME-induced AHT.** (**A**) Experimental design: 10-12-week-old mice were treated with tamoxifen for 5 consecutive days (open arrow heads) and, after 6 weeks, norepinephrine (NE) osmotic minipumps were implanted for 4 weeks in one group of mice; control mice were operated without minipump implantation. Another mouse group was treated with L-NAME in drinking water for 4 weeks. BP was measured at the indicated time points (purple arrows), and mice were euthanized at the end of the experiment. (**B**) Systolic BP values of 11 Cn-Ctl and 6 SM-Cn$^{-/-}$ NE-treated mice, 5 Cn-Ctl and 6 SM-Cn$^{-/-}$ L-NAME-treated mice, and 10 Cn-Ctl and 5 SM-Cn$^{-/-}$ control mice. Data are means ± s.e.m. ***$p<0.001$, **$p<0.01$, *$p<0.05$ vs treated SM-Cn$^{-/-}$, $^{\#\#\#\#}p<0.001$ vs control Cn-Ctl and $^{\#}p<0.05$ vs control SM-Cn$^{-/-}$; RM two-way ANOVA with Tukey's post hoc test.

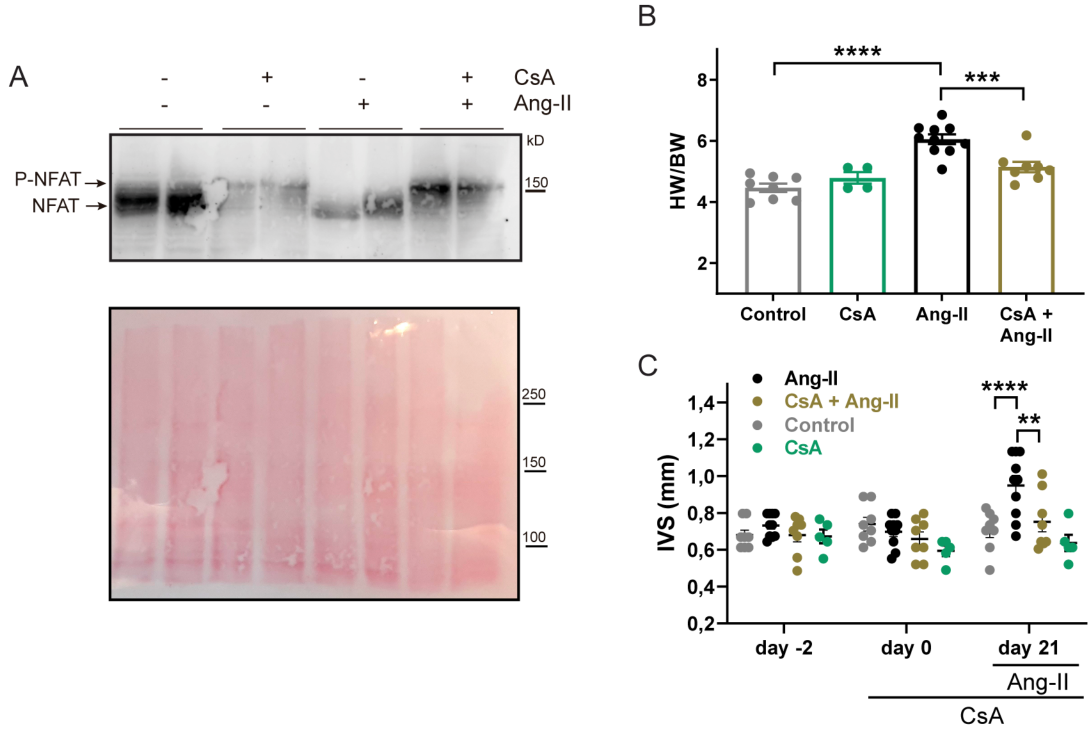


**Supplementary figure 4. Cn inhibition by CsA impairs Ang-II-induced NFAT dephosphorylation and cardiac hypertrophy.** (**A**) Representative NFATc3 immunoblot analysis of thymus extracts from WT mice treated as indicated (top panel) and Ponceau staining of the membrane (bottom panel). Hyper-phosphorylated NFATc3 (P-NFAT) and de-phosphorylated NFATc3 (NFAT) are indicated. (**B**) End-of-experiment heart weight vs body weight ratio (HW/BW) values. Each data point denotes an individual mouse, and data in histograms are presented as mean ± s.e.m. ****$p<0.0001$, ***$p<0.001$; two-way ANOVA with Šídák's post hoc test. (**C**) Echocardiography-determined interventricular septum (IVS) thickness before CsA (day -2) and Ang-II (day 0) administration and after 21 days of Ang-II treatment. Each data point denotes an individual mouse, and the horizontal bars denote the mean (long bar) and the s.e.m. ****$p<0.0001$, **$p<0.01$; RM two-way ANOVA with Tukey's post hoc test. (B-C) Ang-II (n=10), CsA+Ang-II (n=8), control (n=8), and CsA (n=5).

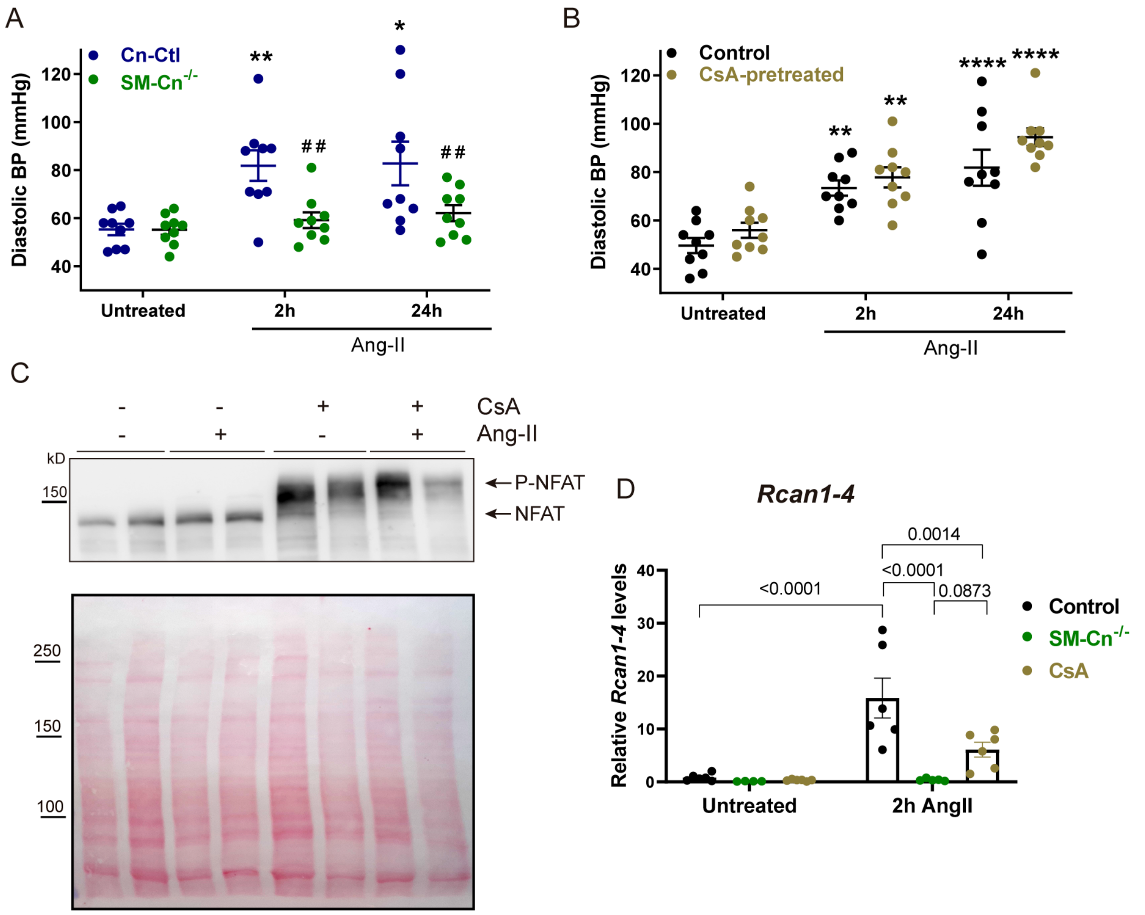


**Supplementary figure 5. Cn deletion, but not inhibition of Cn phosphatase activity, prevents Ang-II-induced diastolic hypertension.** (**A-B**) Diastolic BP was measured at the indicated time points in the same Cn-Ctl, SM-Cn$^{-/-}$, and non-pretreated and CsA-pretreated WT mice as in Figure 6B (A) and Figure 6D (B). Each data point denotes the BP value in an individual mouse, and the horizontal bars denote the mean (long bar) and the s.e.m. (n=9 mice per group and per time point). ****p<0.0001, **p<0.01 vs baseline; RM two-way ANOVA with Tukey's post hoc test. $^{##}$p<0.01, $^{#}$p<0.05 vs 2h or 24h Ang-II Cn-Ctl; RM two-way ANOVA with Šídák's post hoc test. (**C**) Representative NFATc3 immunoblot of thymus extracts from mice treated as indicated. Hyper-phosphorylated NFATc3 (P-NFAT) and de-phosphorylated NFATc3 (NFAT) are indicated. Ponceau staining of the membrane is shown below. (**D**) Quantification of mRNA expression, as assessed by RT-qPCR, in extracts from the aorta of untreated control (3 Cn-Ctl plus 3 WT), SM-Cn$^{-/-}$ (n=5), and CsA-pretreated (n=6) mice and Ang-II-treated control (3 Cn-Ctl plus 3 WT), SM-Cn$^{-/-}$ (n=5), and CsA-pretreated (n=6) mice. Each data point denotes an individual mouse, and data in histograms are presented as mean ± s.e.m. Two-way ANOVA with Šídák's post hoc test, p values, and comparisons are indicated.

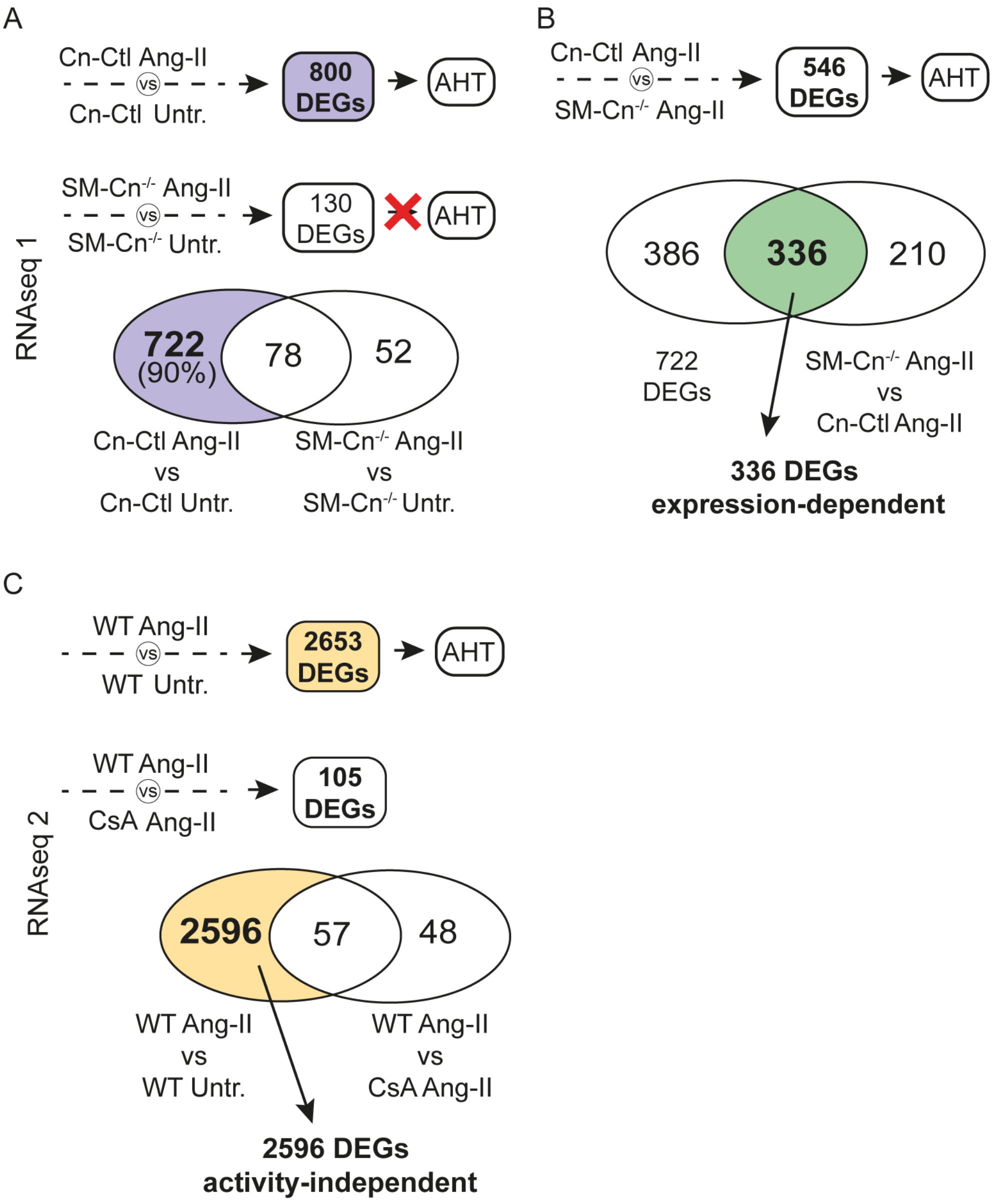


**Supplementary figure 6. Selection of potentially AHT-related genes from the transcriptomic analyses.** (**A**) 800 DEGs in Cn-Ctl mice treated with Ang-II for 2h were selected as potential mediators of AHT, and 130 DEGs regulated in SM-Cn$^{-/-}$ mice treated with Ang-II for 2h were excluded, leaving 722 potentially AHT-related genes, as indicated in the Venn diagram (below). (**B**) Of these genes, only those whose expression differed between the hypertensive condition (Cn-Ctl Ang-II 2h) and the non-hypertensive condition (SM-Cn$^{-/-}$ Ang-II 2h) were selected using a Venn diagram (below). The figure indicates the resulting 336 DEGs potentially involved in AHT and whose regulation is Cn-expression dependent. (**C**) 2653 DEGs regulated in WT mice by treatment with Ang-II for 2h were selected as potential mediators of AHT. Only those whose expression was similar in both hypertensive conditions (not showing differential expression between WT Ang-II 2h and CsA Ang-II 2h) and CsA-independent were selected using a Venn diagram (below). The figure indicates the resulting 2596 DEGs potentially involved in AHT and whose regulation is independent of Cn phosphatase activity.

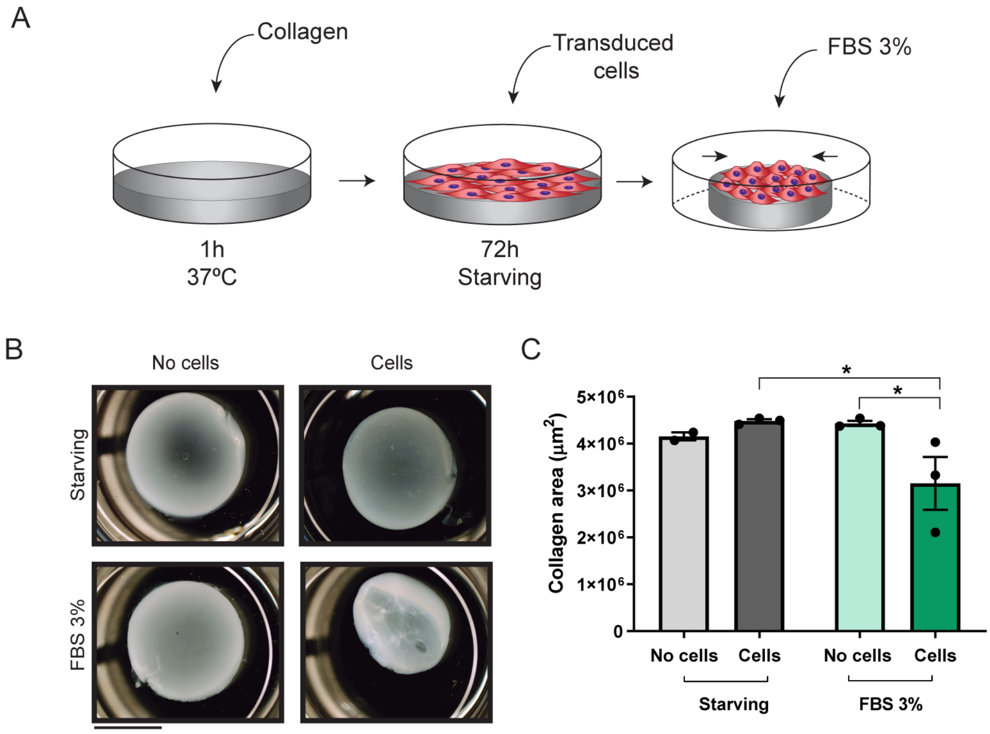


**Supplementary figure 7. Collagen contraction assay.** (**A**) Experimental design: Collagen gels were polymerized for 1h at 37ºC, and VSMCs (8x$10^4$) were seeded on each gel and left to adhere for 5h before being serum-starved for 72h. Gels were then gently detached from the plates and incubated in DMEM supplemented with 3% FBS for 24h-48h at 37ºC. (**B**) Representative images from 3 independent experiments and (**C**) quantification of the surface area of fixed collagen gels in the absence or presence of cells stimulated as indicated. Each data point denotes the mean of each experiment, and data in histograms are presented as mean ± s.e.m. *$p<0.05$ by two-way ANOVA with Šídák's post hoc test. No cells without serum (n=2), starved cells (n=3), No cells with 3% FBS (n=3), Cells with 3% FBS (n=3).
